# Epigenetic progression of pancreatic cancer to aggressive subtypes involves alternate routes of lineage reprogramming in subtype-intermediate progenitor cells

**DOI:** 10.64898/2026.09.28.754674

**Authors:** Kevin A Hawthorne, Jennifer R Eng, Motoyuki Tsuda, Patrick J Worth, Andrew J Fields, Colin J Daniel, Andrew Nishida, Carl Pelz, Ellen M Langer, Trent A Waugh, Ethan S Agritelley, Jason M Link, Brett C Sheppard, Andrew C Adey, Rosalie C Sears

## Abstract

Pancreatic ductal adenocarcinoma (PDAC) progression involves malignant cell state plasticity. Epigenetic changes underlie this plasticity, yet the PDAC *cis*-regulatory landscape remains understudied. To address this, we profiled 33 primary tumors and 7 metastases from 39 patients with single-cell ATAC-seq, paired with 10 single-cell RNA-seq profiles. We found that epigenetic *GATA6*^+^/*KRT17*^+^ co-accessibility identifies a classical-basal subtype-intermediate progenitor state (SIP) associated with better clinical outcomes. SIP cells display limited epigenetic reprogramming from premalignant epithelium and retain gastric-intestinal differentiation reminiscent of neoplastic precursors. Lineages without *GATA6*^+^/*KRT17*^+^ co-accessibility exhibit greater lineage and epithelial-mesenchymal plasticity. Classical PDACs that repress basal gene accessibility activate neural-like progenitor (NRP) and tuft lineage enhancers, whereas basal committed tumors display esophageal transdifferentiation. Compared to SIP, classical-NRP and basal committed tumors have poorer outcomes, and show distinct PD-1/PD-L1 immune proteomic phenotypes and prognostic myofibroblast epigenetic states, respectively. Our work reveals links between lineage reprogramming, EMT, and epigenetic progression in human PDAC.

## Introduction

The past decade of molecular profiling has seen the inclusion of transcriptional subtypes to the molecular taxonomy of PDAC, which have predicted clinical outcome and drug response across numerous independent studies [1–4]. Two consensus subtypes have been repeatedly identified across single-cell resolution datasets: “classical”, which retains expression of endodermal regulators and epithelial identity, and “basal-like”, which is more poorly differentiated and predicts poorer outcome [1, 5]. Spatial and single-cell transcriptomic studies reveal that basal-like and classical cells have different preferences for oncogenic pathways and transcriptional regulators, correlate with distinct stromal and immune compositions, and predict aggressive behavior and metastasis [6–8]. This subtype-specific tumor biology suggests that unique therapeutic vulnerabilities are likely rooted in basal and classical identity. However, knowledge of basal and classical cell states has yet to translate to clinical improvements, and questions remain regarding how these subtypes are regulated and convert between one another.

Cell phenotypes are built not only of the cell’s transcriptome, but also of other critical layers of identity such as the epigenome. Epigenomic features that regulate gene transcription, such as chromatin accessibility, can also encode cell identity and lineage information [9]. For example, overexpression of the proto-oncogene MYC is a near-universal feature of cancer transcriptomes, yet cancer epigenomes vary dramatically by which lineage-specific *cis*-regulatory elements are used to accomplish dynamic MYC dysregulation [10]. This enhancer selection can even affect drug response [11]. Furthermore, transcriptionally similar cells can harbor dissimilar chromatin states [12], and epigenetic memory of prior stressors and metaplastic transitions creates a foundation for malignant cell state plasticity [13, 14]. It is well-known that PDAC subtypes harbor distinct epigenomic landscapes [15–17], but very little is known about this heterogeneity at single-cell resolution, especially in human disease.

Basal-like and classical subtypes display recurring phenotypes and conserved gene transcription programs [18, 19]. These programs emerge in part from the activity of cell fate regulators and lineage specifiers such as TP63 and PDX1 [15, 20]. These observations suggest that PDAC subtypes are encoded in the epigenome as lineage-differentiated states, yet no comprehensive examination of lineage programming has been performed in human PDAC to date.

To determine the epigenetic underpinnings of PDAC molecular subtypes, we profiled 40 human PDAC tumors with single-cell combinatorial indexing (sci)-ATAC-seq. We performed a deep investigation of PDAC subtype programs, uncovering their TF regulatory context, the normal lineages/cell type associated with their enhancers, and their relative deviation from the non-cancer epithelial state. Furthermore, we correlated these findings with epigenetic programs from stromal and immune cells.

## Results

### Single-cell ATAC-seq on human pancreatic tumors

To chart the chromatin accessibility landscape of pancreatic cancer, we performed sci-ATAC-seq on 40 frozen banked tumor disaggregates from 39 patients in the Oregon Pancreas Tissue Registry (OPTR) (previously described in Link JM, et al. [21]). Our sci-ATAC-seq cohort includes 31 primary PDAC tumors, 2 pancreaticobiliary-subtype Ampulla of Vater (PAV) primary tumors suspected to be PDAC in origin, and 7 distant PDAC metastases including liver (n=4), lung (n=2), and ovary (n=1) (Fig. 1a). To navigate data quality challenges inherent to banked pancreatic and tumoral tissue, we imposed rigorous sci-ATAC-seq quality filters (see Methods; Supplementary Fig. 1a-f) to obtain a substantially smaller but refined set of high-quality cells (Fig. 1b). We identified epithelial/malignant, stromal, and immune cell types in primary tumors and metastases (Fig. 1c-d). Lung metastases were enriched for T cells, consistent with prior reports [22, 23], while the ovary metastasis was enriched for stromal cells. Liver metastases uniformly returned few quality cells. 37 of 40 sci-ATAC-seq samples have matched bulk RNA-seq data from our previous publication [21]. In the present work, we provide a supporting scRNA-seq dataset comprising 9 primary PDACs and 1 PAV (Supplementary Fig. 1g-h) samples matched to our sci-ATAC-seq cohort.

**Figure 1:**
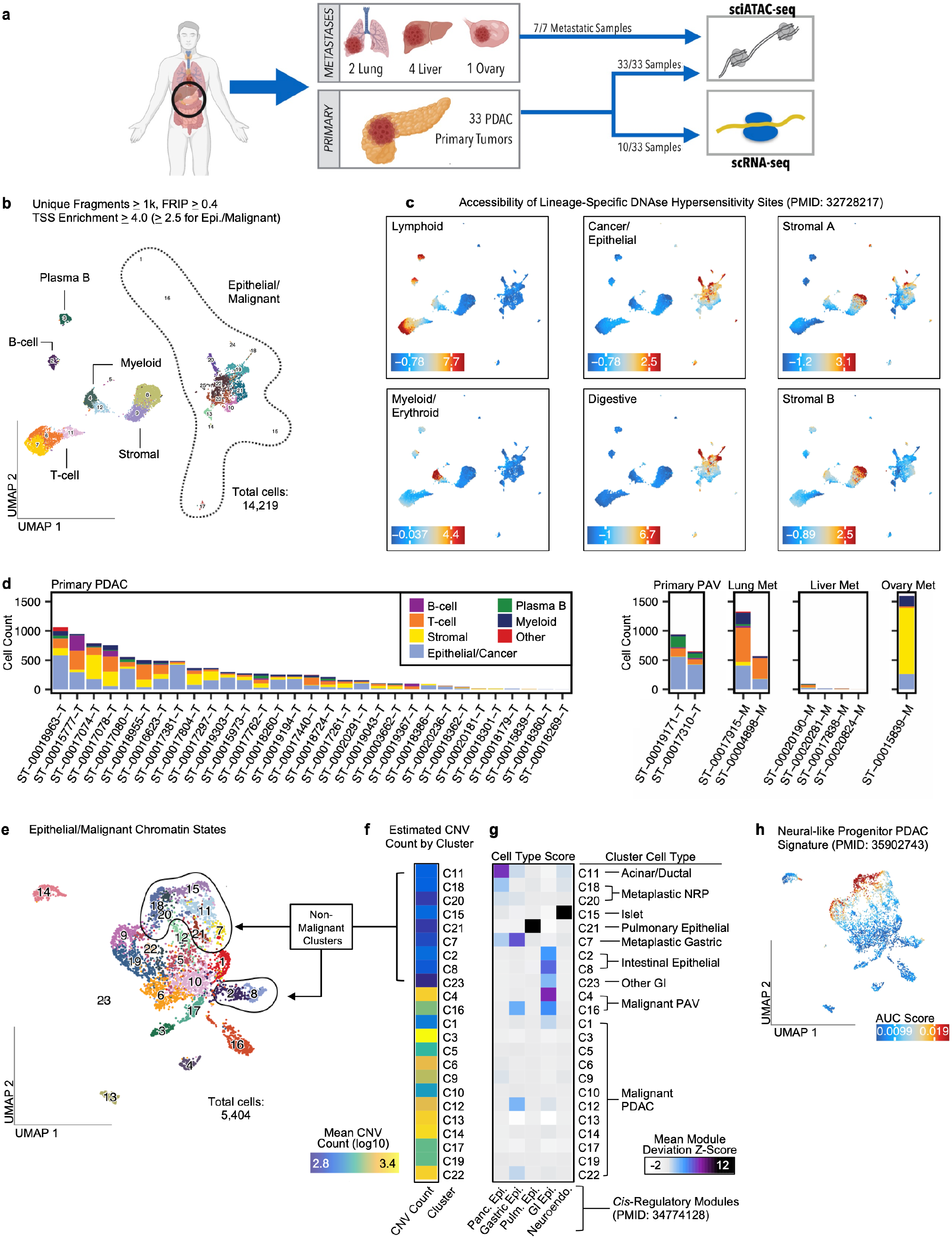
Single cell ATAC-seq captures diverse epithelial, stromal, and immune cell types from PDAC primary tumors and metastases. **a)** Single cell sequencing cohort and sample types. **b)** Single cell clustering results and uniform manifold approximation and projection (UMAP) embedding of all quality-controlled cells from all samples. Each cluster is depicted with a unique cluster and number. All cells have > 1k unique fragments and at least 40% fraction of reads in peaks (FRIP). FRIP is calculated as the proportion of reads overlapping DNase hypersensitivity sites identified by Meuleman W, et al. [72]. Per-cell TSS enrichment is > 2.5 for epithelial/malignant cells, > 4 for stromal and immune. **c)** Per-cell enrichment scores for lineage-associated peak sets by Meuleman W, et al. Enrichment values are bias-corrected deviation Z-scores calculated by chromVAR. **d)** Cell type proportions by sample. The X-axis depicts sample ID; the Y-axis depicts total cell count. Colors represent cell type. **e)** Topic-based single cell clustering and UMAP embedding of all epithelial/malignant cells from all samples (n=5404). Colors and numbers specify individual clusters. Encircled clusters are non-malignant. **f)** Estimated copy number variations (CNVs) by cluster. Row numbers refer to clusters shown in panel a). Values depict the log10 transformation of the average CNV count (gains + losses) of cells in each cluster. **g)** Per-cluster enrichment of lineage-specific regulatory DNA regions from Zhang K and Hocker JD, et al. [25]. Columns depict enrichment results for 5 enhancer signatures selected from the “module” collection from [25]. Row numbers refer to clusters in panel A. Values are the mean per-cell deviation Z-score (via chromVAR) for each cluster. **h)** Enrichment of peaks near genes in the “Neural-like Progenitor” PDAC signature identified by Hwang WL, et al. [7]. Values refer to per-cell Area Under the Curve (AUC) scores calculated by cisTopic. Colors are scaled from the 5th to 95th score percentile.

### Identification of normal, metaplastic, and malignant epithelial chromatin states

To identify epithelial/malignant chromatin states and their underlying regulatory programs, we applied topic modeling via cisTopic [24]. This approach defines co-accessible enhancer programs (topics) and predicts their activity in cells to identify distinct epigenetic states. Topic modeling of epithelial/malignant epigenomes (n=5404) from all samples revealed 50 regulatory topics (Supplementary Fig. 2a) and 23 single-cell clusters (Fig. 1e). We annotated clusters as malignant (n=14) or non-malignant (n=9) manually based on estimated copy number variations (Fig. 1f) (Supplementary Fig. 2b-c). Clusters were generally well-integrated across different patients and tumor anatomical sites (Extended Fig. 1a-d). However, some malignant clusters were patient-specific (Extended Fig. 1d). This may reflect the evolution of distinct, patient-specific subclones that harbor unique copy number aberrations.

We defined cluster cell types based on their accessibility of published human cell type *cis*-regulatory modules [25] (Fig. 1g). Primary PDAC samples contributed acinar and ductal pancreatic epithelial cells, which grouped together into a single cluster (C11; Extended Fig. 1e-f) possibly due to acinar-to-ductal metaplasia (ADM), and pancreatic islets (C15). Lung metastases contributed pulmonary epithelial cells (C21). Intestinal epithelial cells (C2, C8) originated from primary PAV tumors and one primary PDAC (Extended Fig. 1d), likely sampled during tumor resection. Of note, PAV-specific malignant clusters (C4, C16) displayed gastrointestinal (GI) enhancer accessibility on-par with the normal intestinal cells (Fig. 1g). This suggests malignant PAV cells are intestinal-differentiated, despite originating from pancreatobiliary-subtype tumors in our cohort.

We also identified metaplastic cells (C7, C18, C20) based upon two characteristics: normal CNV profile (Fig. 1f), which suggests non-cancer identity, and reduced pancreatic epithelial differentiation (Fig. 1g). We deemed cluster C7 as metaplastic gastric due to its accessibility of gastric epithelial-associated enhancers (Fig. 1g), which suggests gastric-like metaplasia similar to pancreatic pre-cancer models [26, 27]. In contrast, clusters C18 and C20 showed poor accessibility of pancreatic epithelial, gastric epithelial, and islet-associated neuroendocrine modules (Fig. 1g), but displayed accessibility of the neural-like progenitor (NRP) PDAC signature by Hwang WL, et al. [7] (Fig. 1h). This suggests that the PDAC NRP program can occur in non-cancer metaplastic cells, consistent with recent lineage tracing in mouse [28]. We thus refer to C18 and C20 as “metaplastic NRP”.

### *KRT17* and *GATA6* co-accessibility reveals a prevalent subtype-intermediate chromatin state

We next sought to annotate malignant clusters as basal-like or classical, the predominant molecular subtypes of PDAC. Initially, we attempted to discern cluster subtype from accessibility of published basal/classical signatures [2, 3, 7, 29] (Extended Fig. 2a-d). However, basal-like accessibility scores highlighted a substantial proportion of the malignant population (Extended Fig. 2b-d). This was unexpected, as only 12% of malignant cells in our epigenomic dataset originate from tumors deemed by PurIST [30] to be “strong basal-like” by mRNA transcription. Furthermore, classical and basal-like regulatory programs are thought to be mutually opposed [19]. However, subtype scores showed clear overlap in several subpopulations (Extended Fig. 2a-d). This suggested potential epigenetic subtype-intermediate states, reminiscent of basal-classical co-expression in transcriptomic analyses, but much more prevalent [29, 31, 32].

To accommodate subtype-intermediate epigenetic states, we stratified malignant clusters into three epigenetic subtypes based on chromatin accessibility of the classical marker *GATA6* and the basal marker *KRT17* (Fig. 2a-b). Using acinar/ductal cells as a baseline for *GATA6* and *KRT17* accessibility (see Methods; Extended Fig. 2e-f), we classified three malignant clusters as basal *KRT17^+^* (C3, C13, C17), three clusters as classical *GATA6^+^* (C9, C14, C19), and the remaining eight as subtype-intermediate *KRT17^+^ GATA6^+^* co-accessible (Fig. 2c). Of note, two of eight subtype-intermediate clusters originate from primary PAV tumors (C4, C16) which we grouped separately as “Malignant PAV” due to their outlying degree of intestinal differentiation (Fig. 1g).

**Figure 2:**
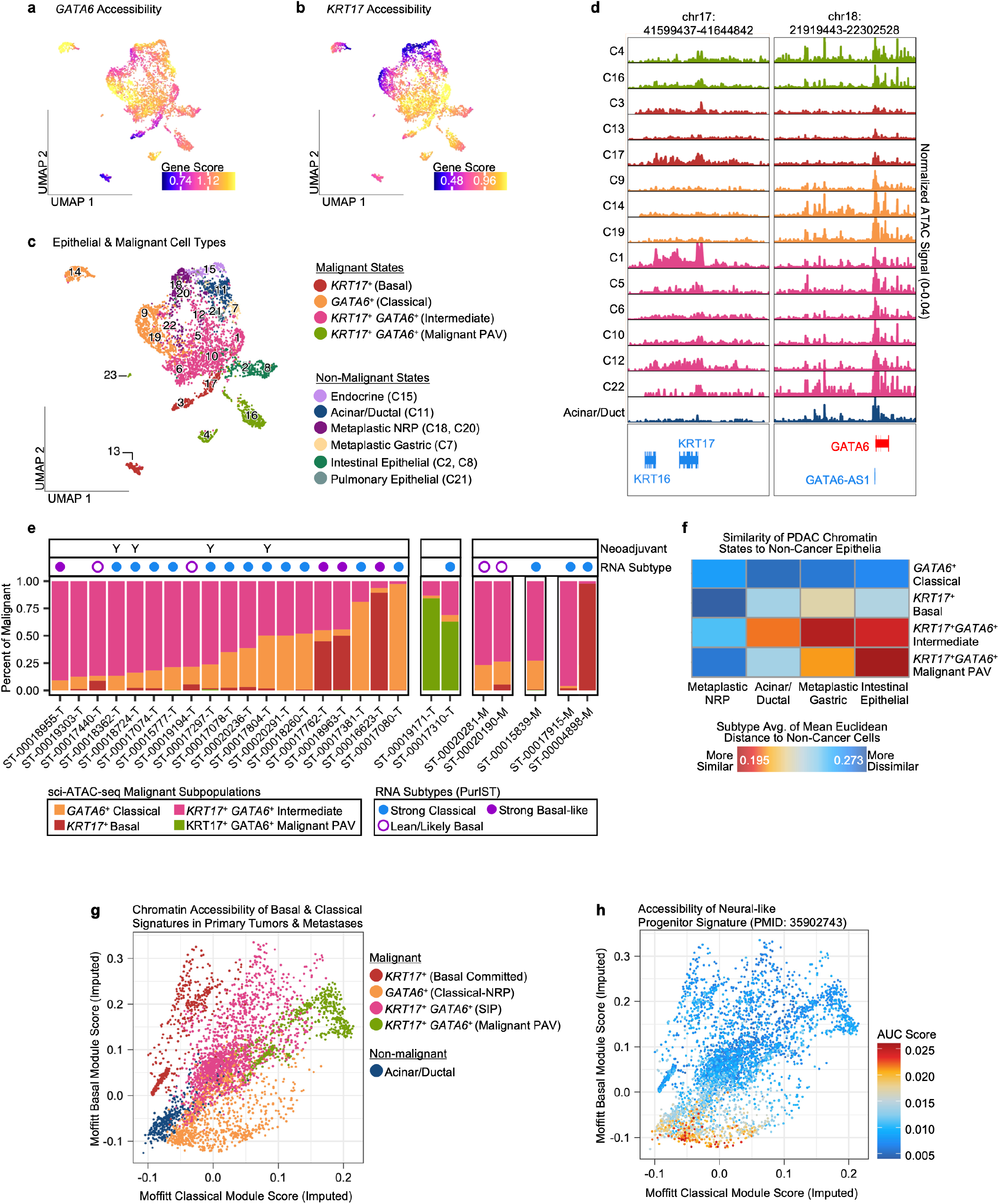
Subtype-intermediate chromatin state is prevalent and precedes clinically unfavorable PDAC subtypes. **a-b)** Imputed *GATA6* and *KRT17* chromatin accessibility scores visualized on UMAP. Colors are scaled from the 0th to 90th score percentile. **c)** UMAP depicting malignant states and non-malignant states. Numbers depict the number and approximate position of the original single cell cluster. **d)** Genomic distribution of ATAC-seq signal near *KRT17* and *GATA6*. Rows depict data for each cluster. Colors correspond to cell type/malignant state as in panel c). **e)** Sample composition by malignant epigenetic subtype. Only samples with greater than 10 malignant cells are shown. **f)** Chromatin state similarity between malignant primary tumor cells and non-cancer epithelial cell types. Color scale depicts the average mean Euclidean distance of malignant cells to a specific group of non-cancer epithelial cells. **g-h)** Per-cell imputed module scores for basal-like and classical gene signatures. In g), cells are colored by annotated subtype. In h), cells are colored by per-cell enrichment of peaks near “Neural-like Progenitor” marker genes. Values refer to per-cell Area Under the Curve (AUC) scores calculated by cisTopic.

Basal *KRT17^+^* malignant clusters repressed putative enhancers upstream of the *GATA6* locus (Fig. 2d). Conversely, *KRT17* was inaccessible in classical *GATA6^+^*clusters, but not basal or subtype-intermediate clusters. Subtype-intermediate clusters showed variable accessibility of other molecular subtype markers, including basal *TP63* and classical *LGALS4* (Extended Fig. 2g-h). In contrast, basal genes *KRT5* and *LY6D* were more restricted to *KRT17^+^* clusters, while classical VSIG2 was most accessible in *GATA6^+^* clusters.

These results suggest intermediate epigenomes have varied levels of access to subtype programs, possibly reflecting the direction of ongoing or future cell state transitions.

Most samples in our sci-ATAC-seq cohort have matched bulk RNA-seq profiles generated in connection to our prior study [21]. sci-ATAC-seq samples that were predominantly basal *KRT17^+^* or classical *GATA6^+^* matched to their bulk PurIST [30] “strong basal-like” and “strong classical” transcriptional profiles, respectively (Fig. 2e). Interestingly, subtype-intermediate-predominant sci-ATAC-seq samples corresponded to transcriptional “strong classical” PurIST subtype; the few that matched to basal-like RNA-seq profiles showed weaker basal transcription (“lean basal-like”, “likely basal-like”). Although the epigenetic subtype-intermediate PDAC cells have basal-like gene accessibility, these results suggest that the intermediate chromatin state nevertheless produces one-sided transcription in favor of the classical subtype.

Previous spatial proteomic and transcriptomic profiling indicate that subtype-intermediate “co-expressor” cells are present in most PDAC tumors, but generally as a minor subpopulation [29, 33, 34]. In contrast, the *KRT17^+^ GATA6^+^* co-accessible intermediate chromatin state was prevalent in our sci-ATAC-seq cohort (Fig. 2e). While most tumors harbored a predominant epigenetic subtype, the three epigenetic subtypes co-existed to varying extents, indicating intra-tumoral heterogeneity. Importantly, although many tumors were split between two major epigenetic subtypes, this was always between one subtype and the *KRT17^+^ GATA6^+^* co-accessible intermediate; no tumor was split primarily between basal *KRT17^+^* and classical *GATA6^+^* cells. Taken together with the “strong classical” gene expression of epigenetic intermediate PDAC tumors, these results suggest widespread chromatin accessibility, but not gene transcription, of the basal-like program among malignant cells.

### Subtype-intermediate state is a progenitor to advanced malignant subtypes

Recent observations raise the possibility that the basal-classical intermediate expression state and *KRT17* gene expression itself can be achieved early in disease progression [35–37]. Given the prevalence of subtype-intermediate epigenomes in our dataset, we investigated if the intermediate *GATA6^+^ KRT17^+^* state precedes the *GATA6^+^* state, the *KRT17^+^*state, or both. To do so, we determined the chromatin state similarity of each subtype to non-cancer epithelial cell types in primary tumors (Fig. 2f). Subtype-intermediate cell epigenomes showed shorter Euclidean distances to normal and metaplastic cell types compared to the basal *KRT17^+^* and classical *GATA6^+^* states. This suggests that the *GATA6^+^ KRT17^+^* co-accessible intermediate cell state is less reprogrammed from the pre-cancer state, and thus likely precedes the *KRT17^+^*and *GATA6^+^* accessible subtypes. We therefore refer to this subtype as “subtype-intermediate progenitor” or “SIP” moving forward.

Classical PDAC is thought to retain more endoderm-pancreatic identity and regulation than basal-like PDAC [1, 38]. However, *GATA6^+^* cell epigenomes displayed greater deviation from acinar/ductal chromatin state than the basal *KRT17^+^* cells (Fig. 2f), suggesting they comprise a deeply reprogrammed yet non-basal subtype. A subset of the *GATA6^+^* population displayed NRP [7] program activation concomitant with loss of the Moffitt RA, et al. [3] classical program (Fig. 2g-h). This suggests *GATA6^+^* epigenomes shift to an NRP phenotype, a clinically unfavorable and drug-persistent cell state [7]. Because *GATA6^+^* epigenomes activate the NRP program, yet originate from transcriptionally classical tumors (Fig. 2e), we refer to this group as the “classical-NRP” or “C-NRP” subtype. In contrast, we refer to *KRT17^+^*epigenomes as the “basal committed” or “BC” subtype to reflect their singular activation of the basal-like program.

### *Cis*-regulatory programs associated with PDAC epigenetic subtypes

To characterize the SIP, classical-NRP, and basal committed epigenetic subtypes, we leveraged the results of our earlier topic model, which identified 50 topics/regulatory programs active in epithelial and malignant cells (Extended Fig. 3a). Our linear mixed-effects modeling (LMM) approach (see Methods) identified 19 subtype-associated topics that showed significantly higher scores (LMM contrast FDR<0.05) in one malignant subtype compared to the other two across primary PDAC tumors (Fig. 3a). We analyzed these 19 *cis*-regulatory programs for enrichment of 1) enhancers near canonical transcriptional subtype genes (Fig. 3b), 2) TF binding motifs for a curated list of pancreatic progenitor [20], exocrine, endocrine, and EMT regulators (Fig. 3c), and 3) accessible regions for 222 human cell types from a published atlas [25] (Fig. 3d-e). The results of these analyses, detailed for each epigenetic subtype in the following sections, unveil the TF regulation and lineage differentiation programs underlying PDAC subtype heterogeneity.

**Figure 3:**
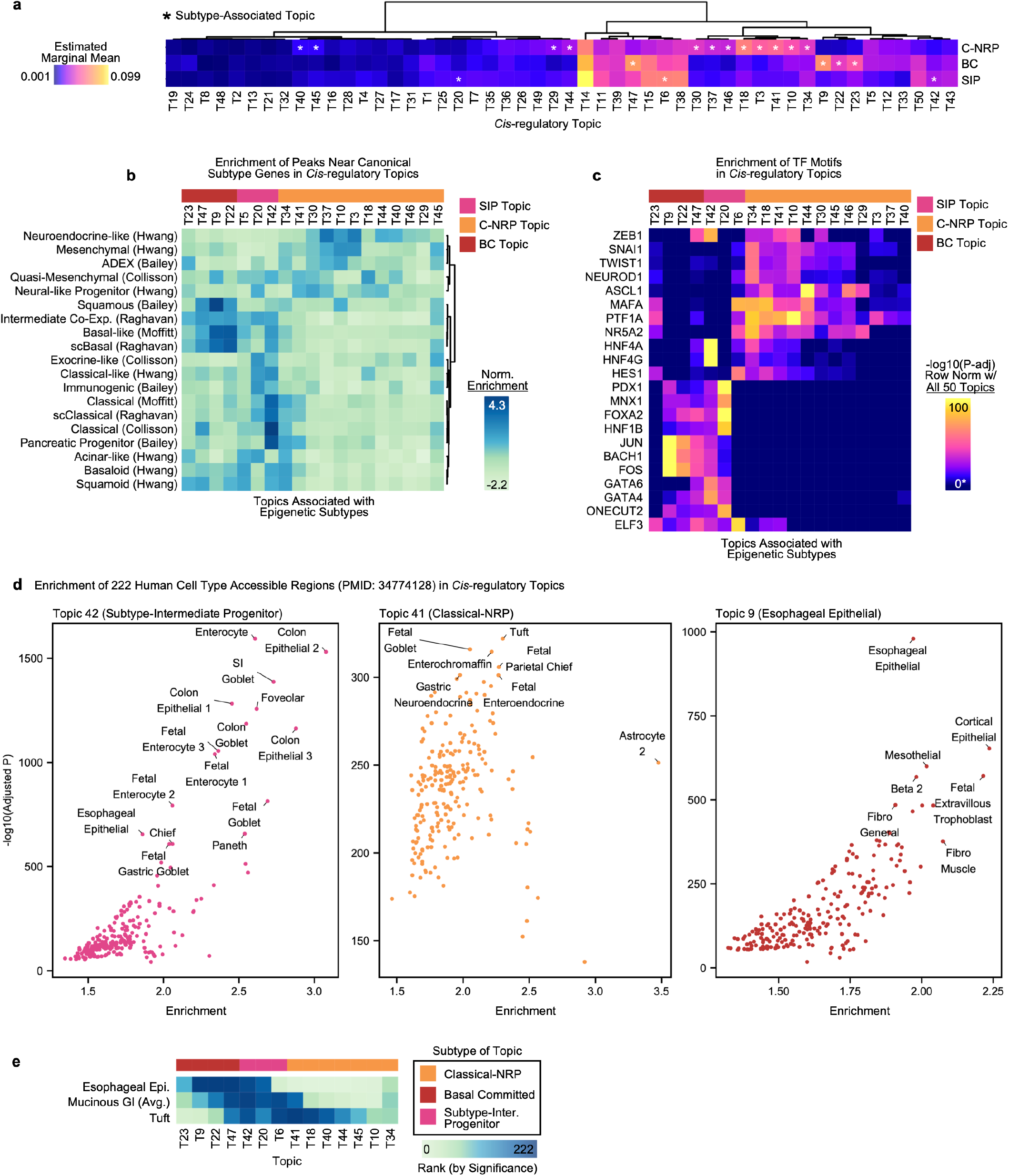
PDAC *cis*-regulatory programs reveal three distinct axes of malignant cell differentiation. **a)** Identification of subtype-associated regulatory topics in primary PDACs (excluding PAV tumors). Colors depict marginal means estimated from our LMM workflow (see Methods). Asterisks indicate topics that showed significantly greater marginal means (LMM contrast FDR<0.05) in one subtype compared to the other two subtypes. FDR correction was applied to contrasts within each LMM (by topic) **b)** Per-topic enrichment of peaks near genes in published PDAC subtype signatures. Color scale depicts normalized enrichment scores calculated by cisTopic. The subtype that was associated with each topic (from panel a) is depicted by the top color bar. Rows were clustered via Ward method using correlation distances. **c)** Per-topic hypergeometric enrichment of TF motifs. Colors depict-log10 hypergeometric adjusted p-values that were row-normalized (0-100) across all 50 topics in our model; only the 19 subtype-associated topics are shown. * Non-significant (adjusted P < 0.05) results were scaled to the 0 value on the color scale. Bonferroni correction was applied to P-values across all tests (870 motifs tested per topic * 50 topics total). **d)** Per-topic hypergeometric enrichment of cell type enhancers in example regulatory topics. Each point represents a cell type peak set (222 total) collected from Zhang K and Hocker JD, et al. [145]. Colors refer to the epigenetic subtype associated with the topic. Bonferroni correction was applied to P-values across all tests (222 cell types tested per topic * 50 topics total). **e)** Significance rank by p-value of representative cell type peak sets across subtype-associated topics. “Mucinous GI” depicts the average rank of goblet, foveolar, enterocyte, and colon epithelial peak sets in a given topic. 5 classical-NRP topics were excluded due to having extremely few (<5) significant results.

### Subtype-intermediate PDAC retains PanIN-like differentiation and progenitor-like TF regulation

All regulatory programs associated with subtype-intermediate progenitor (Topics 6, 20, 42) showed enrichment (NES>0.5) of the “intermediate co-expressor” signature by Raghavan S et al. [29] (Fig. 3b). Of the three SIP programs, basal enhancers and classical enhancers co-occurred specifically in Topic 42, which enriched (NES>0.5) nearly all basal-type and classical-type signatures in our analysis. TF motifs significantly enriched (p<0.05, Bonferroni-adjusted) in Topic 42 included AP-1 factors similar to basal committed programs (e.g. Topic 9), but also GATA6 and GATA4, which regulate canonical classical phenotype [39] (Fig. 3c). All SIP topics showed enrichment of various TF motifs corresponding to PDAC pancreatic progenitor TFs identified by Bailey P, et al. [38]. These include significant enrichment (p<0.05, Bonferroni-adjusted) of HES1 (Topic 6); HNF1A/B, FOXA2/3, PDX1, and MNX1 (Topic 20); and HNF4A/G (Topic 42) (Fig. 3c). These results underscore that SIP is a progenitor-like identity, driven by endoderm cell fate TFs that regulate pancreatic development and early PDAC progression.

Consistent with our earlier finding that SIP resembles non-cancer states (Fig. 2f), SIP-associated topics were also accessible in normal epithelial and metaplastic gastric cells (Extended Fig. 3b-d). SIP programs also highlighted the metaplasia-associated TF ONECUT2 [40], both through high TF motif enrichment (enrichment score (ES)=2.03, Bonferroni-adjusted p=6.72×10^-85^ in Topic 20; Fig. 3c) and by encompassing accessible regions near the *ONECUT2* gene locus (Extended Fig. 3e). Furthermore, SIP topics showed high TF motif enrichment of ELF3 (ES=2.09, Bonferroni-adjusted p=1.31×10^-94^ in Topic 6; Fig. 3c), which enforces epithelial identity and represses EMT [41], and consistent de-enrichment (NES<-0.5) of the mesenchymal PDAC signature by Hwang WL et al. [7] (Topics 6, 20, 42; Fig. 3b). These results suggest SIP retains partial accessibility of pre-malignant enhancers, and maintains a potentially less invasive epithelial-adherent phenotype.

SIP lineage infidelity was characterized by gastric and intestinal epithelial cell types. Topic 42 (Fig. 3d, left) and Topic 6 (Extended Fig. 4a) showed significant enrichment (Bonferroni-adjusted p<0.05) of mucin-expressing GI lineages including enterocyte, foveolar, and goblet cells. Topic 20 co-enriched gastric chief and pancreatic acinar/ductal cell enhancers (Bonferroni-adjusted p<0.05; Extended Fig. 4b). Gastric-like differentiation states are prototypical of pancreatic neoplasia and emerge during pre-malignant PDAC development [26, 27]. Our results indicate SIP is programmed predominantly as a gastric-intestinal epithelial identity, which is reminiscent of gastric-like neoplastic precursors that emerge during early tumor development.

Tuft and esophageal epithelial lineages were associated with classical-NRP and basal committed programs, respectively (Fig. 3d). Although the top ranking lineages in SIP topics were mucinous GI lineages (mean p-value rank between 202.8 to 219.0; 222 max), tuft and esophageal epithelial lineages also ranked highly (mean tuft rank in SIP topics=190.7, esophageal epithelial=140.0) (Fig. 3d). This may potentially reflect epigenetic priming of other subtype programs by SIP cells, similar to multilineage priming by progenitor cells in normal development.

### Classical-NRP reveals a tuft-neuroendocrine differentiation axis in PDAC

Classical-NRP was associated with a high number of regulatory programs (n=12 topics), reflecting complex cell state transitions along the classical-to-NRP trajectory. Although some classical-NRP programs showed enrichment of multiple canonical classical signatures (NES>0.5; Topics 34, 41), the majority (8 of 12) enriched the NRP and/or NE-like programs [7] (Fig. 3b). Five of the classical-NRP epigenetic programs also showed enrichment of the “aberrantly differentiated endocrine exocrine” (ADEX) subtype signature by Bailey P et al. [38], which was often co-enriched (NES>0.5) with the mesenchymal PDAC signature [7]. Accordingly, TF motifs for endocrine fate regulators (ASCL1, MAFA, NEUROD1, REST), exocrine fate regulators (PTF1A, NR5A2) and EMT regulators (TWIST1, SNAI1-2, and ZEB1) were all co-enriched (Bonferroni-adjusted p<0.05) in classical-NRP topics (Fig. 3c). These results suggest that the classical-NRP identity is EMT-competent and co-regulated by endocrine and exocrine TFs.

Lineage analysis revealed that classical-NRP cells were accessible for tuft and neuroendocrine enhancers. Topic 41, the top classical-NRP program of the 12 by mean score (Extended Fig. 3a), showed pronounced enrichment (ES>1.98, Bonferroni-adjusted p<0.05) of tuft and several GI neuroendocrine cell types (Fig. 3d, middle). In contrast, the top 2^nd^ and 3^rd^ programs (Topics 18, 10) highlighted neuronal/glial cell types (Extended Fig. 4c-d), which likely reflects cross-expression of neural markers by NRP cells [28] rather than neuronal differentiation. Furthermore, both classical-NRP and metaplastic NRP cells shared Topic 41 activation and accessibility of the tuft marker *DCLK1* (Extended Fig. 4e-f). Our results suggest that the NRP program is common to both malignant and metaplastic cells in human PDAC and involves activation of tuft and NE enhancers, consistent with the tuft-to-NRP differentiation trajectory recently elucidated in mouse [28]. These data support the existence of a tuft-neuroendocrine differentiation axis in human PDAC, which likely exists within the broad umbrella of the canonical classical subtype.

### Basal committed PDAC shows AP-1 regulation and esophageal lineage infidelity

Regulatory programs associated with the basal-committed epigenetic subtype (Topics 9, 22, 23, 47) strongly aligned with canonical basal gene expression signatures (Fig. 3b). Befitting the most aggressive PDAC subtype, basal-committed topics contained numerous peaks near genes involved in oncogenic signaling pathways. These include epidermal growth factor signaling (*EGFR*), MAPK signaling (*ERK1/MAPK3*, *ERK2/MAPK1*), and multiple oncogenes (e.g. *KRAS, AXL, AKT2*). Basal-committed programs showed enrichment of TF motifs for AP-1 (JUN, FOS, BACH1; Fig. 3c), which mediates epigenetic responses to inflammation and oncogenic KRAS [42, 43]. We also noted peaks proximal to circadian factors *NPAS2*, *ARNTL*, and *BMAL2/ARNTL2* (Extended Fig. 5a-b). In our primary tumor bulk RNA-seq dataset (n=218), *BMAL2* mRNA expression was greater in basal-like PDACs compared to classical (p=3.5×10^-10^), correlated with *HIF1A* expression (R=0.31, p=2.2×10^-6^), and predicted poorer clinical outcome (p=7.3×10^-4^) (Extended Fig. 5c-e). These results suggest basal-like regulation is sustained by AP-1 and circadian-related factors in PDAC.

The basal committed subtype displayed a shift from pancreatic to esophageal identity, evidenced by the striking enrichment of esophageal epithelial enhancers (ES>1.97, Bonferroni-adjusted p<0.05) across topics of this subtype (Fig. 3d; Extended Fig. 5f-g). The esophagus is composed of basal and squamous cell types that naturally express basal-like PDAC markers [44, 45]. Although basal/squamous PDAC has been reported to downregulate genes that maintain endodermal identity [38], our results suggest PDAC basal differentiation is directed towards an endodermal lineage (esophageal).

### All PDAC subtypes access enhancers associated with MYC regulation and prior GI differentiation

We next sought to identify “consensus” peaks that are ubiquitously accessible or silenced across the 3 epigenetic malignant subtypes (SIP, C-NRP, BC) in primary PDAC tumors (excluding PAV tumors). Pairwise differential accessibility tests between acinar/ductal cells and each malignant subtype yielded intersected results of 4.2k consensus PDAC peaks and 571 consensus acinar/ductal peaks (Fig. 4a; differential peaks defined as log2 fold change>1, FDR<0.01).

**Figure 4:**
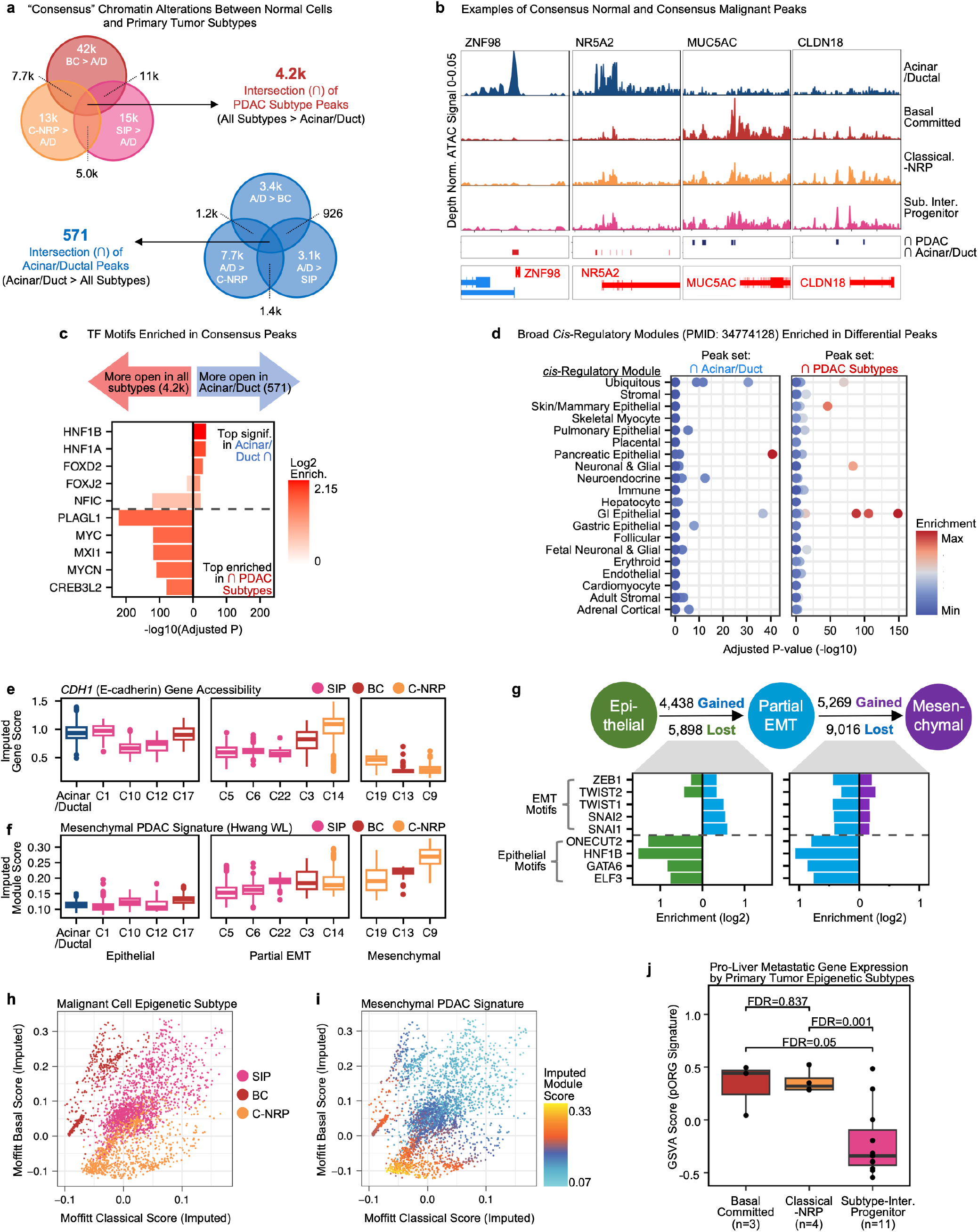
All subtypes retain neoplasia-associated features and MYC regulation, but differ in invasive phenotypes. **a)** Differentially accessible peaks (binomial FDR<0.01, log2 fold change>1) identified between each epigenetic subtype and acinar/ductal cells from primary PDAC tumors. Top Venn diagram depicts peaks more accessible in each subtype, bottom diagram depicts peaks more accessible in acinar/ductal cells. **b)** Genomic distribution of ATAC-seq signal near *ZNF98*, *NR5A2*, *MUC5AC*, and *CLDN18*. Rows depict ATAC-seq signal from different cell types from primary PDAC tumors. Box below cell type rows depicts chromatin regions identified from panel a). Genes depicted in the bottom box are red if they are encoded on the plus strand, blue if encoded on the minus strand. **c)** Top TF motifs significantly enriched (hypergeometric adjusted p<0.05) in consensus PDAC peaks (left boxplot) and acinar/ductal peaks (right boxplot). Bonferroni p-adjustment was applied across all tests (2 peak sets * 870 TF motifs). Negative log2 enrichment scores are scaled to 0. **d)** Lineage-specific *cis*-regulatory modules enriched in consensus acinar/ductal peaks and consensus PDAC peaks (hypergeometric adjusted p<0.05). Enrichment scores in acinar/ductal peaks range from 0 (min) to 23.3 (max); scores in PDAC peaks is range from 0 (min) to 5.53 (max). Each point represents a module peak set collected from Zhang K and Hocker JD, et al. [25]. Modules are organized by their associated lineage (rows). Bonferroni p-adjustment was applied across all tests (2 peak sets * 150 modules). **e)** Imputed *CDH1* gene accessibility score in acinar/ductal cells and malignant clusters (primary PDAC tumors and metastases). Clusters are colored by their malignant subtype (SIP, BC, C-NRP) and grouped by their EMT state (epithelial, partial EMT, mesenchymal). **f)** Imputed gene signature module scores for the mesenchymal PDAC signature by Hwang WL, et al.[7]. **g)** *Top:* Count of differentially accessible peaks (binomial FDR<0.01, log2 fold change>1) between epithelial cells and partial EMT cells, and partial EMT cells and mesenchymal cells. *Bottom:* Hypergeometric enrichment of TF motifs in differential peaks. Green bars indicate TF motifs enriched in epithelial peaks, blue bars indicate motifs enriched in partial EMT peaks, and purple bars indicate motifs enriched in mesenchymal peaks. **h)** Epigenetic subtype of malignant cells, plotted against their per-cell imputed module scores for the Moffitt RA basal and classical signatures (X and Y axes). **i)** Per-cell imputed module scores for the mesenchymal PDAC signature (same data as panel f) plotted against classical and basal module scores (same as panel h). **j)** pORG score of bulk RNA-seq profiles matched to primary PDAC samples in the sci-ATAC-seq cohort. Scores were compared via Wilcox test.

Among genes near consensus PDAC peaks, we noted genes involved in fundamental oncogenic processes, including cell cycle dysregulation (*MYC*), telomere maintenance (*TERT*, *TEP1*), PP2A suppression (*PPP2R2D*), and proteolytic growth factor release (*KLK* family) (Extended Fig. 6-7a-d). In contrast, multiple consensus acinar/ductal peaks (n=8) occurred near the acinar differentiation TF *NR5A2*, suggesting PDACs frequently repress this locus (Fig. 4b). The primate-specific TF *ZNF98* was silenced in virtually all malignant cells, suggesting it may function as an uncharacterized PDAC tumor suppressor. TF motifs enriched in consensus acinar/ductal peaks featured HNF1A/B and NFIC (Fig. 4c), which regulate pancreas development and acinar identity [46–48]. In contrast, the top enriched TF motifs in consensus PDAC peaks corresponded to MYC and related factors (MAX, MXI1, MYCN), underscoring the central role of MYC in PDAC development and progression [49] (Fig. 4c). These results suggest consensus PDAC peaks encompass core tumorigenic pathways, whereas consensus acinar/ductal peaks encompass TFs that maintain pancreatic lineage and function.

Among genes near consensus PDAC peaks, we noted markers of PanIN precursor lesions, including *MUC5AC* [50, 51], *CLDN18* [50, 51], and *CTSE* [52] (Fig. 4b; Extended Fig. 7e). In addition, consensus PDAC peaks showed significant overlap with GI epithelial enhancers (Fig. 4d). These results imply that all 3 epigenetic subtypes can emerge from a prior mucinous epithelial state, and epigenetic memory of this previous identity is partially retained. Gastric and intestinal epithelium have naturally high cell turnover; this memory may confer partial GI characteristics that favor growth and survival. Consistent with this, consensus PDAC peaks upstream of the pro-mitotic oncogene *MYC* overlapped normal GI lineage enhancers (Extended Fig. 6).

### Basal committed and classical-NRP subtypes associate with EMT and pro-liver metastatic gene expression

Our earlier topic analysis suggested that SIP is an epithelial identity, whereas classical-NRP is mesenchymal. To investigate this, we compared malignant clusters (see Fig. 2c) by their chromatin accessibility scores for E-cadherin and the mesenchymal PDAC signature by Hwang WL et al. [7], also incorporating metastatic cells to understand the full range of invasive phenotypes (Fig. 4e-f). We classified malignant clusters with high E-cadherin score, yet low mesenchymal scores, as epithelial (C1, 10, 12, 17). We labeled three clusters (C9, 13, 19) as fully mesenchymal due to their high mesenchymal scores and near complete loss of E-cadherin signal. The remaining clusters (C3, 5, 6, 14) were deemed “partial EMT”. Classical-NRP showed only partial EMT (C14) and mesenchymal (C9, C19) states (Fig. 4e-f), suggesting this subtype leans towards an invasive phenotype. In contrast, basal committed clusters covered the entire EMT spectrum, from epithelial (C17) to partial EMT (C3) to mesenchymal (C13). SIP, despite comprising 45% of total malignant cells in our model, did not form a fully mesenchymal *CDH1^-^* cluster. This suggests that SIP cells are unable to complete EMT, or that EMT occurs jointly with differentiation into basal committed or classical-NRP subtype.

Next, we investigated how EMT states differ by regulation. Differential accessibility tests between EMT states (irrespective of subtype) indicated that the shift from epithelial state to partial EMT was accompanied by opening of ZEB, TWIST, and SNAI-regulated peaks, and closure of peaks recognized by pro-epithelial TFs ELF3, HNF1B, GATA6, and ONECUT2 (Fig. 4g, left barplot). Upon shifting from partial EMT to mesenchymal state, peaks regulated by epithelial TFs and EMT TFs alike both appear to close (Fig. 4g, right barplot), while new EMT TF recognition sites open. These results suggest that peaks recognized by pro-epithelial TFs close progressively over the course of EMT, whereas recognition sites for EMT TFs appear to change throughout the process.

Visualizing per-cell mesenchymal gene signature module scores against basal/classical module scores depicts a putative progression of basal committed and classical-NRP states to EMT (Fig. 4h-i). In this scheme, the classical-NRP trajectory to EMT is defined by progressive silencing of the Moffitt RA et al. [3] classical program, concomitant with activation of mesenchymal genes [7]. Classical-NRP cells with the highest mesenchymal module scores show negative module scores for the Moffitt RA classical program (Fig. 4i). This suggests an EMT trajectory that is inextricably related to subtype identity, as classical-NRP cells must completely repress the Moffitt RA classical program to accomplish a completely mesenchymal chromatin state. In contrast, basal committed cells progressing to mesenchymal state show reduced (but still positive) module scores for the Moffitt RA basal program (Fig. 4i). This suggests basal PDAC does not need to substantially modify its lineage differentiation state to switch between epithelial-mesenchymal phenotypes.

Our analyses suggest basal committed and classical-NRP are metastatic phenotypes due to their EMT proficiency, whereas SIP may be less invasive. To investigate this, we examined if epigenetic subtypes show different mRNA expression of our published primary organotropism (pORG) signature [21]. High pORG indicates expression of genes associated with primary tumors that develop liver metastatic disease. Tumors with high pORG also show high replication stress, suppressed T-cell responses, and poor clinical outcome [21]. RNA-seq profiles for basal committed and classical-NRP primary tumors in our sci-ATAC-seq cohort showed significantly greater pORG scores than SIP (FDR<0.05; Fig. 4j). This suggests that classical-NRP and basal committed epigenetic subtypes are associated with prognostically unfavorable, pro-liver metastatic gene expression.

### Fibroblasts from SIP tumors resemble normal stroma

Next, we investigated if PDAC epigenetic subtypes produce distinct stromal chromatin states. Topic modeling of stromal epigenomes (n=1957) from primary tumors identified 27 regulatory topics (Extended Fig. 8a) and 8 stromal cell clusters (Fig. 5a). We identified myofibroblastic CAFs (myCAFs; clusters C1, C2), inflammatory CAFs (iCAFs; C3, C4), and pancreatic stellate cells (PSCs; C5, C6) based on accessibility of PDAC stromal gene signatures [3, 6, 53] (Fig. 5b-d). We combined iCAF clusters and PSC clusters respectively (Fig. 5e), as marker peak analysis revealed minimal intra-cluster differences within these cell types (Extended Fig. 8b). In contrast, myCAF clusters C1 and C2 showed dissimilar marker peak profiles (Fig. 5f), indicating they are distinct regulatory states. We labeled C1 as *LRRC15*^+^ myCAFs and C2 as *NGFR*^+^ myCAFs due to their significant marker peaks (FDR<0.05, log2 fold change>0.5) near these genes (Fig. 5e,g). *LRRC15* is a marker of tumor-promoting myCAFs [54, 55], whereas *NGFR*/*CD271* is expressed by activated stellate cells [56] and early tumor-reactive PSCs [57]. Antigen presenting CAFs (apCAFs) have also been reported in PDAC [58], but we did not observe CAFs with accessibility for MHC-II loci in our dataset (Extended Fig. 8c).

**Figure 5:**
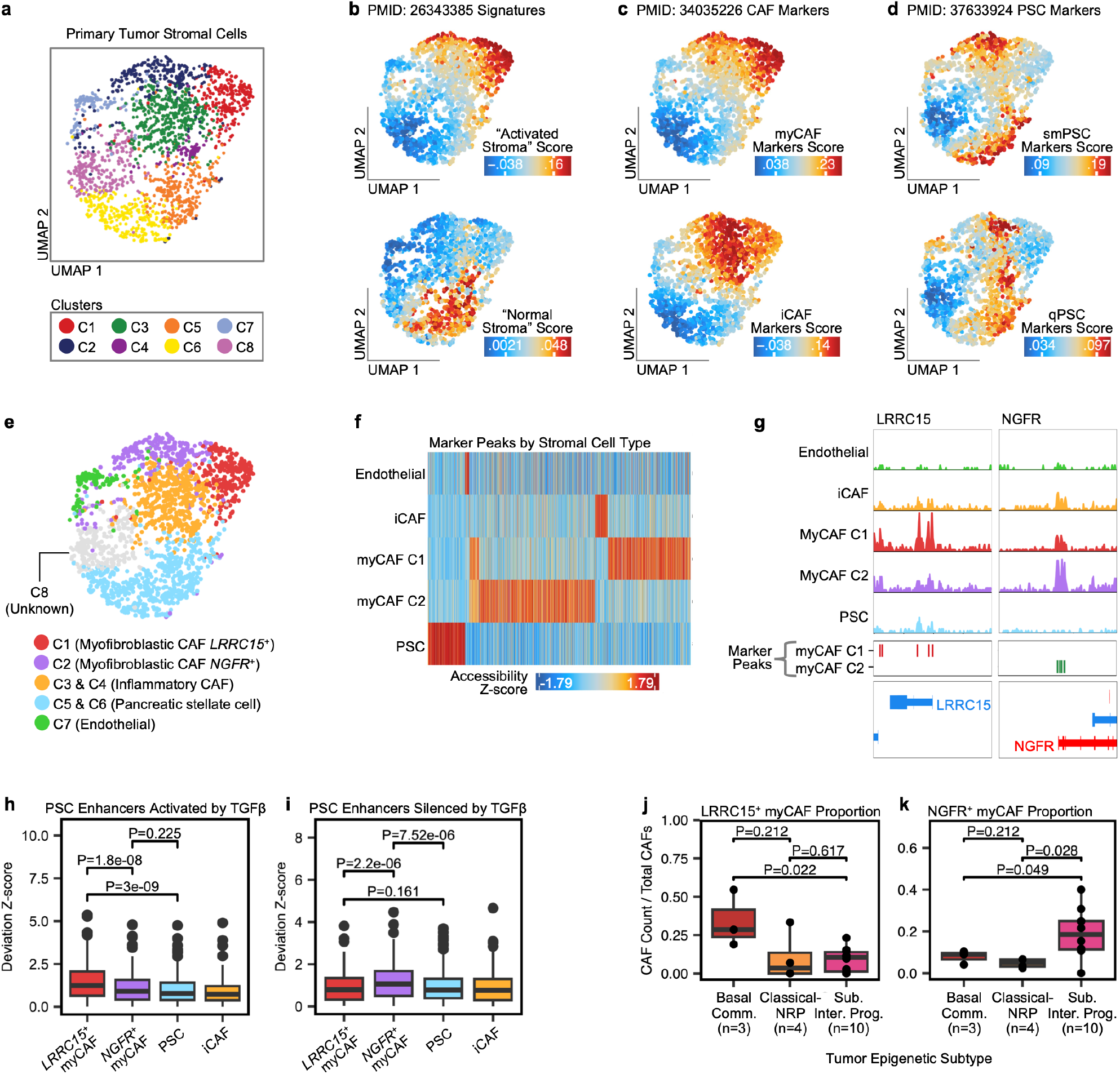
MyCAFs from SIP PDAC tumors retain more normal, stellate-like chromatin accessibility. **a)** UMAPs of stromal cell epigenomes from primary tumors. Colors depict cell clusters. **b-d)** UMAPs depicting per-cell imputed module scores for stromal/CAF gene signatures from published studies. **e)** UMAP annotated by stromal cell type and CAF subtype. Clusters (e.g. C1) refer to clustering shown in panel a). **f)** Heatmap of significant marker peaks (binomial FDR<0.05 & log2 fold change>0.5) identified for each stromal cell type (rows) by ArchR. Columns depict individual peaks. Color scale represents the Z-scored accessibility of the peak. **g)** Genomic distribution of ATAC-seq signal near TSS regions for *LRRC15* and *NGFR.* Rows in the top box depict the ATAC-seq signal from each cell type indicated. The middle box depicts the location of significant marker peaks (from panel f) for C1 myCAFs and C2 myCAFs. Bottom box depicts the genomic location of genes; red genes are encoded on the plus strand, blue on minus. **h-i)** Per-cell chromatin accessibility of TGFβ-activated and TGFβ-repressed PSC enhancers. Scores are deviation Z-scores calculated by chromVAR. Significance was assessed via Wilcox test. **j-k)** Proportion of *LRRC15^+^* myCAFs and *NGFR^+^* myCAFs out of total CAFs in primary PDAC tumors. Significance was assessed via Wilcox test.

*NGFR^+^* myCAFs (C2) showed accessibility of PSC-related gene sets (Fig. 5d), suggesting a PSC-like phenotype. PSCs are a source of CAFs in PDAC [59] and differentiate into myofibroblasts in response to TGF-β [60]. *LRRC15^+^* myCAFs are also known to be TGF-β-driven [54, 55]. To determine if *NGFR^+^*and *LRRC15^+^* myCAF states are both regulated by TGF-β, we analyzed a separate sci-ATAC-seq dataset profiling TGF-β response in human PSCs (see Methods) [61]. PSC enhancers activated by TGF-β showed greater accessibility in *LRRC15^+^* myCAFs (p<1.8×10^-8^; Fig. 5h). In contrast, PSC enhancers silenced by TGF-β showed more accessibility in *NGFR^+^* myCAFs (p<7.52×10^-6^; Fig. 5i). This suggests that *NGFR^+^* myCAFs retain enhancers associated with the pre-activated PSC state compared to *LRRC15^+^* myCAFs, which show stronger TGF-β-related chromatin remodeling. Importantly, by proportion of total CAFs, *LRRC15^+^*myCAFs were enriched in basal committed PDACs compared to SIP (p=0.022), whereas *NGFR^+^* myCAFs were enriched in SIP tumors overall (p<0.049; Fig. 5j-k). Tumor subtype did not affect PSC or iCAF proportions (Extended Fig. 8d-e). Together, these results suggest myCAFs from SIP tumors retain more PSC-like chromatin accessibility, whereas myCAFs from basal committed tumors are TGF-β-polarized to the *LRRC15^+^*state.

### Regulatory programs underlying CAF subtypes reveal novel transcriptional regulators

*LRRC15*^+^ myCAFs showed specific activation of two regulatory programs (Topics 15, 25) (Fig. 6a), which were both enriched for peaks near myCAF markers (Fig. 6b). Gene Ontology (GO) enrichment analysis of Topic 15 revealed skeletogenic processes related to osteoblast function and cartilage development (Fig. 6c). Supporting this, TF motifs for osteoblastic RUNX family members were significantly enriched (FDR<0.05) in Topic 15, in addition to AP-1 factors (Fig. 6d-e). *LRRC15*^+^ myCAFs also activated peaks near *RUNX2*, the master regulator of osteoblast differentiation [52] (Fig. 6f). In our scRNA-seq data, *RUNX2* and its target *DLX5* were expressed by *LRRC15^+^*myCAFs (Extended Fig. 8f-h). These results suggest RUNX factors regulate the *LRRC15^+^* myCAF chromatin state.

**Figure 6:**
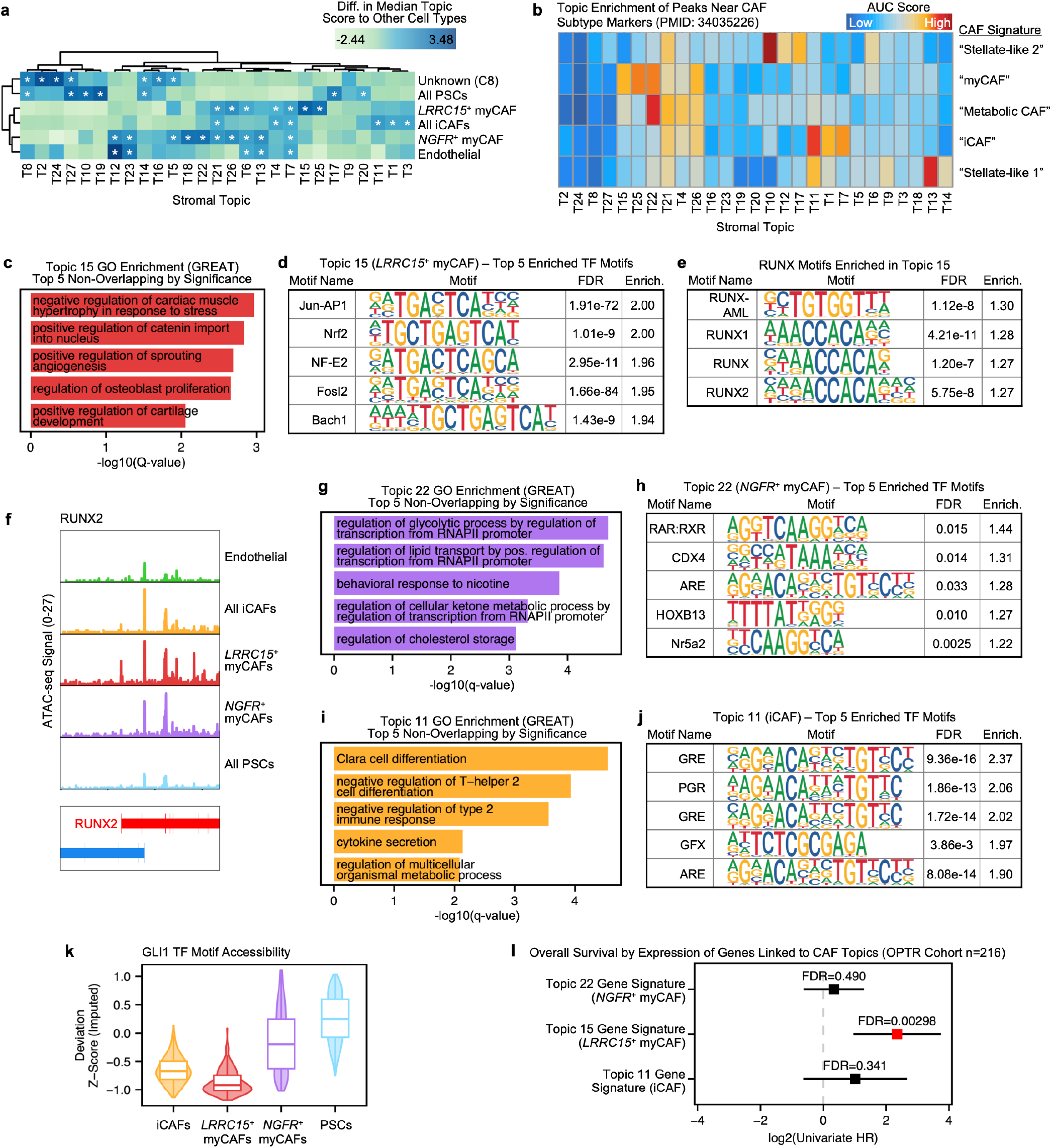
N*G*FR^+^ myCAFs and *LRRC15*^+^ myCAFs are associated with distinct TF regulators and clinical outcomes. **a)** Identification of *cis*-regulatory topics associated with each stromal subpopulation. Asterisks indicate topics that were significantly enriched (T-test FDR<0.05, median log2 fold change>0.5) in a given cell type population compared to all other cells. Color scale depicts the difference between the median score of the row cell type compared to all other cells. **b)** Per-topic enrichment of peaks near CAF subtype genes identified by Wang Y, et al [4]. Color scale depicts the normalized enrichment score calculated by cisTopic. **c)** Top 5 most significant Gene Ontology (GO) Biological Process terms enriched in Topic 15, an *LRRC15*^+^ myCAF program. Biological Process terms with only a single annotated gene are excluded. Redundant terms that are immediately related (in the GO hierarchy) to a term shown in the figure are excluded. **d)** Top 5 most enriched TF motifs in Topic 15 identified by HOMER. P-values were FDR-corrected across all TF motif tests (27 stromal topics * 439 TF motifs). **e)** Significantly enriched RUNX TF motifs in Topic 15 from the same analysis as panel d). **f)** Genomic distribution of ATAC-seq signal near the *RUNX2* locus. Each row depicts the ATAC-seq signal for the labeled stromal cell type. **g-h)** Same as panel c-d) but for Topic 22, an *NGFR*^+^ myCAF program. **i-j)** Same as panel c-d) but for Topic 11, an iCAF program. **k)** Imputed per-cell chromVAR scores for the GLI1 TF motif. **l)** Correlation between gene expression (GSVA scores) of epigenetic CAF signatures and overall survival (as univariate hazard ratios). Significance was assessed via Wald test.

**Figure 7:**
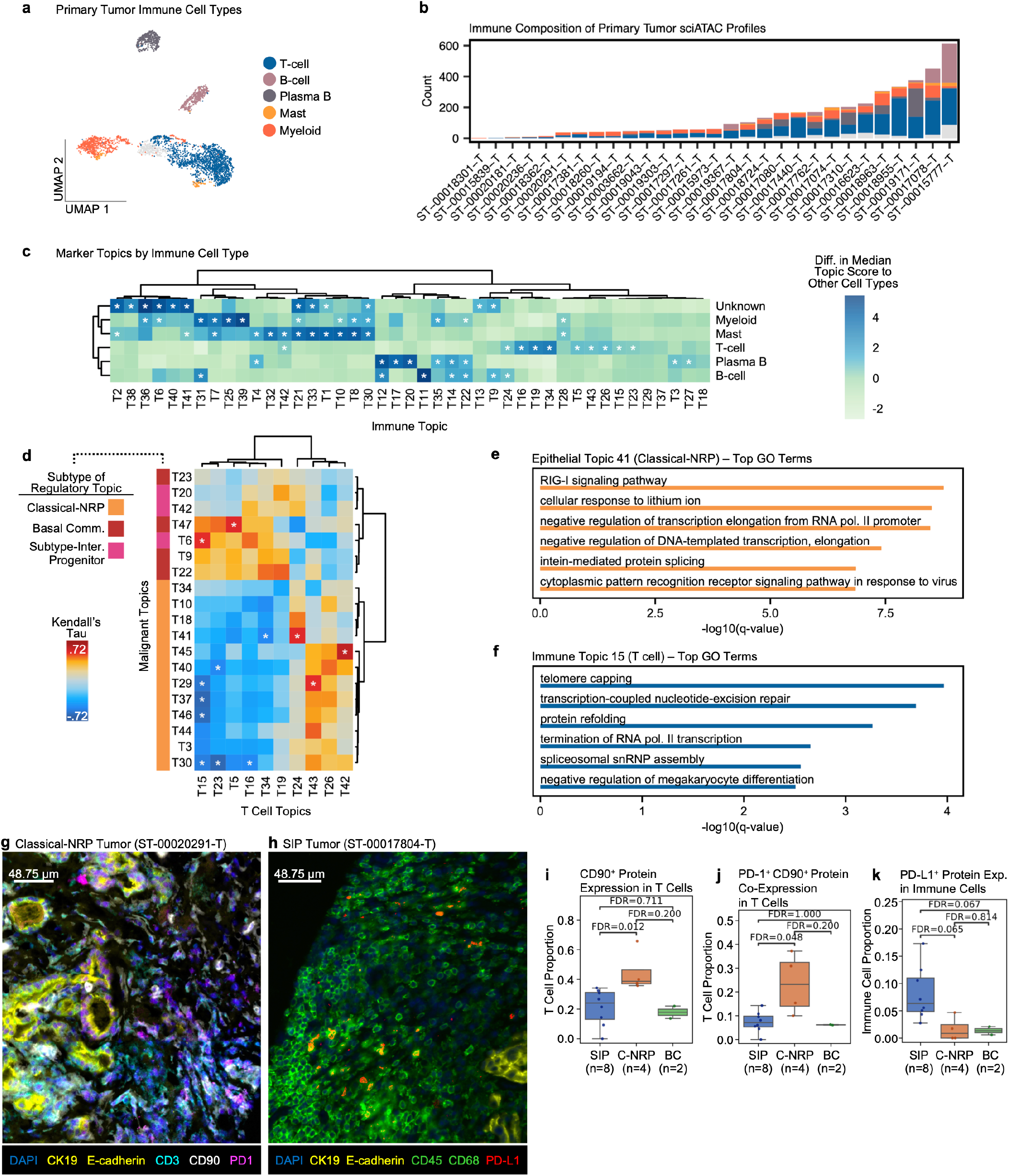
PDAC epigenetic subtypes relate to distinct T cell programs and PD-1/PD-L1 immune phenotypes. **a)** UMAP depicting immune cell types in primary tumors. **b)** Proportion of immune cell types per primary tumor. Colors depict the same immune cell types as in panel A. **c)** Identification of *cis*-regulatory topics associated with immune cell types. Asterisks indicate topics that were significantly enriched (T-test FDR<0.05, median log2 fold change>0.5) in a given cell type population compared to all other cells. Color scale depicts the difference between the median score of the row cell type compared to all other cells. **d)** Sample-level correlation between T-cell associated immune topics (columns) and malignant subtype-associated topics (rows). Asterisks indicate significant (FDR<0.05; Kendall’s Tau) correlations. Color scale depicts correlation value. Both columns and rows are clustered using Ward method under Euclidean distance. **e-f)** Top enriched GO terms from epithelial Topic 41 (classical-NRP) and immune Topic 15 (T cell) peaks determined via GREAT [81]. **g-h)** Representative tumor sections cyclically immunostained for markers depicted at the bottom of each image. **i-j)** Cyclic immunofluorescence analysis of primary PDAC tumor histologic sections. The Y axis depicts the proportion of T cells that show CD90 staining (panel i) or co-staining of CD90 and PD-1 (panel j). Significance was determined via Mann-Whitney U test. **k)** Same as i-j, but depicts the proportion of immune cells that show PD-L1 staining.

*NGFR^+^* myCAFs uniquely activated a regulatory program (Topic 22; Fig. 6a) that contained peaks near myCAF markers, but also peaks near markers of a “metabolic CAF” subtype proposed by Wang Y et al. [53] (Fig. 6b). Consistent with this, GO terms enriched in Topic 22 indicated metabolic regulation and lipid transport (Fig. 6g). The top enriched TF motif in Topic 22 was retinoid X receptor alpha (RXRA) (ES=1.44, FDR=0.015), a nuclear receptor involved in retinoic acid signaling (Fig. 6h). Retinoic acid signaling suppresses the fibrotic program [62]; this result may indicate *NGFR^+^* myCAFs are receptive to pro-quiescence TME signals.

The iCAF-specific regulatory program Topic 11 (Fig. 6a-b) showed enrichment (q<0.05) of immune-modulating GO terms including cytokine secretion (Fig. 6i). Topic 11 displayed high TF motif enrichment (ES>1.90, FDR<0.05) of androgen receptor (AR) and other steroid hormone nuclear receptors (GRE, PGR) (Fig. 6j).

CAFs showed strong AR gene expression in our scRNA-seq dataset, although this was not specific to iCAFs (Extended Fig. 8i). This possibly suggests that AR is expressed in most CAFs, but is signaling actively only in iCAFs.

### *NGFR^+^* myCAFs are not a tumor-promoting CAF subtype

Though *LRRC15^+^* myCAFs are tumor promoting [54, 55], hedgehog-responsive myCAFs potentially restrain metastasis [63, 64]. *NGFR^+^* myCAFs displayed high accessibility of the TF motif for GLI1 (Fig. 6k), a canonical effector of hedgehog signaling. This suggested that *NGFR^+^* myCAFs may restrain tumor progression. To investigate this, we developed stromal-specific transcriptional signatures for each epigenetic CAF subtype (see Methods; Supplementary Fig. 3). In our bulk RNA-seq cohort (n=216 primary tumors), GSVA scores for the *LRRC15^+^*myCAF signature predicted poorer outcome (univariate HR=5.09, Wald FDR=0.00298), consistent with prior characterization [55] (Fig. 6l). In contrast, the *NGFR^+^* myCAF (FDR=0.490) and iCAF (FDR=0.341) signatures were not prognostic. This indicates that the *NGFR^+^* myCAFs are not a tumor-promoting CAF subtype.

### PDAC epigenetic subtypes correlate with distinct T cell programs

Next, we investigated the relationship between immune cell states and PDAC epigenetic subtypes. Topic modeling of primary tumor immune cells (n=4073) yielded 43 regulatory programs and 13 immune cell clusters (C1-C13) (Extended Fig. 9a-b). Marker gene accessibility (Extended Fig. 9c-f) revealed cluster identities as myeloid (C1, C2), B cell (C9, C10), plasma B (C6, C12), and T cell (C3-C5, C13) (Fig. 7a). Marker peaks identified between the remaining unknown clusters (C7, C8, C11) and T cells revealed C7 as putative mast cells and C8 as CD8^+^ T cells, with C11 remaining unknown (Extended Fig. 9g-h). Our dataset captured predominantly CD8^+^ T subsets, as clusters showed T-specific accessibility near *CD3* and *CD8A/B* but not *CD4* (Extended Fig. 9i). T cells were most prevalent (46%), followed by myeloid (18%), plasma B (13%), B cells (12%), and mast (8%) (Fig. 7b).

Tumor epigenetic subtypes did not affect cluster proportions within any immune cell type (Supplementary Fig. 4), suggesting that subtype-related immune reprogramming is unrelated to our clustering results. To uncover changes in regulation, we correlated immune topics by cell type (Fig. 7c) against malignant subtype topics (see Fig. 3a) in primary PDACs (excluding PAV tumors). Mean subtype topic scores within malignant cells (per sample) correlated significantly (Kendall’s Tau FDR<0.05) with mean immune topic scores within T cells (Fig. 7d) but not other immune cell types (Supplementary Fig. 5). These results suggest that PDAC epigenetic subtypes are associated with distinct T cell epigenetic programs. Moreover, T cell topics that correlated positively with classical-NRP topics showed negative correlations with SIP and basal committed topics and vice-versa (Fig. 7d). This suggests T cells from classical-NRP tumors have distinct chromatin states relative to SIP and BC.

To explore the relationship between classical-NRP subtype and T cell states, we examined epigenetic regulatory topics in more detail. GO enrichment of the top classical-NRP topic, Topic 41, revealed viral DNA/RNA response terms including the RIG-I pathway (Fig. 7e). This suggests a link between the classical-NRP subtype and innate immune signaling, which promotes production of interferons and pro-inflammatory cytokines. Classical-NRP cells also potentially relate to altered T cell phenotypes due to their tuft-like differentiation (Fig. 3d). Intestinal tuft cells were reported to resist CD8^+^ T-killing despite expressing antigen presentation machinery, possibly by dampening CD8 T proliferation [65]. Relatedly, we observed a putative T cell proliferation stress topic, Topic 15, which showed enrichment of GO terms related to genomic integrity and protein refolding (Fig. 7f). T cell Topic 15 correlated negatively with multiple classical-NRP programs (Topics 29, 30, 37, 46) but positively with SIP Topic 6 (Fig. 7d), suggesting T cells are non-dividing or exhausted in classical-NRP but not SIP. Overall, these results support that PDAC epigenetic subtypes relate to distinct T cell chromatin states.

### Immune cell PD-1 and PD-L1 phenotypes differ between classical-NRP and SIP tumors

To investigate T cell phenotypes at the protein level, we analyzed immune markers in histologic sections of SIP (n=8), basal committed (n=2), and classical-NRP (n=4) primary tumors (Fig. 7g-h; Extended Fig. 10a-b). Cyclic immunofluorescence staining of T cell markers did not reveal differences in CD4^+^ T or FOXP3^+^ T between PDAC epigenetic subtypes (Extended Fig. 10c-d). However, classical-NRP tumors were significantly enriched for CD90^+^ T cells compared to SIP (Fig. 7i; p=0.004, FDR=0.012). T cells that co-expressed CD90 and the inhibitory receptor PD-1 (CD90^+^ PD-1^+^) were also enriched in classical-NRP compared to SIP (Fig. 7j; p=0.016, FDR=0.048), whereas single-positive PD-1^+^ T cells were not enriched in any subtype (Extended Fig. 10e).

These results suggest that CD90^+^ PD-1^+^ T cells represent a subset of T cells associated with the epigenetic classical-NRP subtype that have seen chronic activation and are exhausted. In contrast, immune cells that expressed the PD-1 ligand PD-L1 showed striking enrichment in SIP tumors (Fig. 7k) compared to classical-NRP (p=0.022, FDR=0.065) and basal committed (p=0.044, FDR=0.067). These results suggest that the classical-NRP and SIP epigenetic subtypes are associated with distinct PD-1 and PD-L1 immune phenotypes.

### Subtype-intermediate PDAC features correlate with favorable outcome

Our analyses overall suggest SIP tumors are less dysregulated across multiple tumoral compartments. We therefore hypothesized that SIP tumors have better clinical outcome. Within our sci-ATAC-seq cohort, patients with SIP primary tumors (n=12) survived longer than patients with classical-NRP (n=4; log-rank FDR=0.049) or basal committed tumors (n=3; log-rank FDR=0.042) (Fig. 8a). This contrasts with a previous model that places “intermediary” gene expression subtype outcomes in-between basal-like and classical [34]. Additionally, because SIP and classical-NRP both relate to classical gene expression (Fig. 3e), this result suggests that multiple clinically distinct epigenetic states exist within the canonical classical PDAC umbrella.

**Figure 8:**
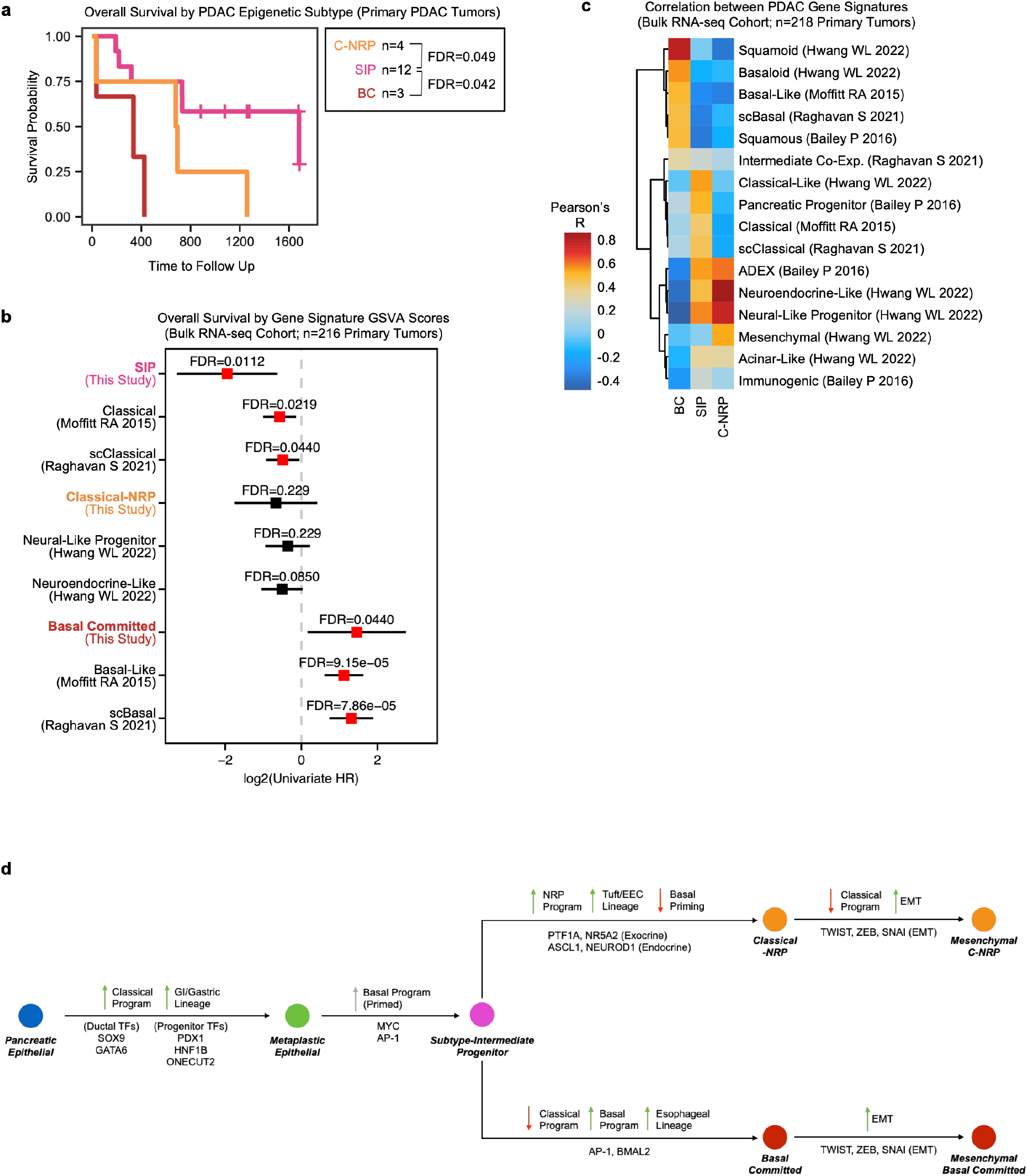
SIP is a clinically favorable epigenetic subtype. **a)** Overall survival of primary PDACs in the sci-ATAC-seq cohort, stratified by the predominant malignant subtype within each PDAC tumor. Significance assessed via log-rank test. **b)** Univariate hazard ratio based upon gene expression (GSVA score) of published gene signatures and sci-ATAC-seq gene signatures from this study. Significance assessed via Wald test. **c)** Pearson correlation between published gene signatures and sci-ATAC-seq gene signatures from this study. Correlation is calculated from gene expression GSVA scores. **d)** Putative hierarchy of malignant differentiation states in PDAC. Pancreatic epithelial cells may transition to canonical classical/mucinous GI identity upon neoplastic transformation. The basal program is presumed to be epigenetically primed at this stage, or upon PDAC initiation as the subtype-intermediate progenitor. The progenitor state then differentiates towards classical-NRP or basal lineages.

Next, we investigated mRNA expression of genes near SIP enhancers and their relationship to clinical outcome. We identified 84 genes near 127 SIP-specific regulatory sites (see Methods), which we refer to as the “SIP signature”. In our scRNA-seq dataset, gene expression of the SIP signature overlapped both ductal-like cells and KRAS-mutant cells (Supplementary Fig. 6a-c). In our larger bulk RNA-seq cohort (n=216), SIP signature expression predicted lower risk of death (univariate HR=0.259, Wald FDR=0.0112) and had a lower hazard ratio compared to canonical classical PDAC signatures [3, 29] (Fig. 8b). The SIP signature shares few genes with classical signatures (2 on average) [2, 3, 7, 29, 38]. Despite this, SIP signature expression correlated predominantly with classical signatures in the OPTR cohort (Fig. 8c).

We also developed classical-NRP and basal committed signatures from our epigenetic subtype-specific regulatory regions (n=200 genes each; see Methods). Expression of the basal committed signature predicted greater risk of death (univariate HR=2.74, Wald FDR=0.0440) and correlated with published basal signatures (Fig. 8b-c). In contrast, the classical-NRP signature correlated with NRP, NE, and mesenchymal PDAC gene sets [7], consistent with our epigenetic analysis (Fig. 3b). Neither our classical-NRP signature nor the original NRP signature by Hwang WL affected risk of death (Fig. 8b). However, this may reflect cross-expression of NRP markers by non-malignant cells, as remarked in the original study [7].

Overall, our survival analysis suggests that the epigenetic classical-NRP and basal committed PDACs have worse outcomes than SIP. From this result and our other sci-ATAC-seq characterization (Fig. 2-4), we hypothesize that PDAC epigenetic progression initiates with the progenitor-like SIP and advances to the more invasive classical-NRP or basal committed subtypes through closing of enhancers for the basal or classical program, respectively, and progressive lineage infidelity involving opening of alternate cell lineage states (Fig. 8d).

## Discussion

While much is understood about PDAC transcriptional subtypes, this knowledge has been difficult to translate to clinical benefits, and little is known about the epigenetic basis from which these subtypes emerge. By profiling human PDAC tumors with single-cell ATAC-seq, our study reveals that the basal and classical subtype programs are co-accessible in a subtype intermediate, progenitor-like malignant state (SIP) whose epigenetic phenotypes resemble that expected of early disease. SIP PDAC cells show limited pathological chromatin remodeling, epithelial-mesenchymal plasticity, and lineage infidelity compared to other epigenetic subtypes. Furthermore, CAFs from SIP-predominant PDAC tumors exhibit more normal, stellate-like chromatin accessibility. These results suggest that the epigenetic SIP subtype comprises a prognostically favorable subgroup of PDAC tumors that are characterized by limited epigenetic progression and stromal reprogramming.

Our analyses uncovered an EMT-competent subgroup of transcriptionally classical PDAC tumors: the epigenetic classical-NRP subtype. This group showed accessibility of the NRP program recently identified by Hwang WL, et al. [7]. The NRP subtype expresses neuronal markers such as NRXN3 and was linked to poor outcome in the original study [7]. In our work, cells within the classical-NRP group showed an identity shift from the gastric-intestinal epithelial identity (SIP) to the NRP identity. The origin of malignant NRP cells in human PDAC has been uncertain due to cross-expression of NRP markers by nonmalignant epithelium. In mice, a recent lineage tracing model by Salas-Escabillas DJ, et al. revealed that metaplastic tuft cells undergo neuroendocrine differentiation to NRP cells in the progression to carcinoma [28]. Classical-NRP cells and metaplastic NRP cells share *cis*-regulatory topics in our dataset, suggesting that both cell states actuate a common differentiation program. These topics were enriched for tuft cell enhancers; thus, our data suggest that the tuft-to-NRP differentiation trajectory modeled in mouse [28] operates in human pancreatic disease, both in non-malignant and malignant cells. Furthermore, TF motifs enriched in classical-NRP topics suggest that the NRP state is co-regulated by endocrine and exocrine TFs, similar to the ADEX subtype reported by Bailey P, et al. [38]. The ADEX subtype was previously thought to represent contamination [66]; our data suggest ADEX is a regulatory state underlying classical-NRP.

We found that SIP, classical-NRP, and basal committed epigenetic subtypes share a core set of GI epithelial enhancers. This is consistent with the notion that PDACs can emerge from a prior GI-like or mucin-expressing PanIN/IPMN precursor [26, 27]. Our results suggest SIPs do not substantially modify their differentiation state relative to that expected of neoplastic precursors. In contrast, basal committed and classical-NRP subtypes diverge into alternate endodermal fates: esophageal epithelial in the former, and tuft/enteroendocrine in the latter. Esophageal lineage infidelity supports the prediction that PDAC traverses the shared esophagus-pancreas endoderm developmental path to achieve basal differentiation [67].

Each epigenetic subtype was associated with specific stromal or immune epigenetic states. Basal committed PDACs were enriched for the epigenetic *LRRC15^+^* myCAF state. SIP was associated with the more stellate-like *NGFR^+^* myCAF state; this state showed high GLI1 TF motif accessibility and may thus relate to hedgehog-responsive, anti-tumor myCAFs [64]. In the immune compartment, PDAC epigenetic subtypes were associated with distinct T cell programs and phenotypes. Notably, classical-NRP tumors were enriched for a subset of T cells that co-express CD90 and the inhibitory receptor PD-1, which may reflect chronic antigen exposure, inflammation, or other TME factors that promote T cell exhaustion. On the other hand, SIP tumors were uniquely enriched in PD-L1^+^ immune cells. This indicates that the classical-NRP and SIP epigenetic subtypes, which both correspond to PurIST “strong classical” gene transcription [30] (Fig. 2e), may have dissimilar responses to immunotherapy. These results support the well-understood role of molecular subtypes in modifying other tissue compartments in PDAC.

Our work also produced salient findings regarding the role of malignant subtypes and tumor progression. SIP was associated with better clinical outcome and poor epithelial-mesenchymal plasticity. In contrast, both the classical-NRP and basal committed subtypes displayed competence for a complete mesenchymal transition. They were also associated with worse clinical outcome than SIP. These results suggest two paths to a mesenchymal identity: basal commitment, which involves esophageal epithelial differentiation; or classical-NRP, which involves tuft-neuroendocrine differentiation. Apart from EMT, the common link between the classical-NRP and basal committed subtypes is their severe degree of epigenetic reprogramming and lineage infidelity. Both subtypes showed greater dissimilarity to non-cancer epithelial states than SIP, suggesting that they both undergo more substantial chromatin remodeling.

## Methods

### Tissue acquisition and patient consent

Banked human tissue samples for single-cell sequencing datasets described in this work were acquired through the Oregon Pancreas Tissue Registry (OPTR) under the OHSU Institutional Review Board (IRB) study #3609. Analysis of molecular data collected from these human tissues was performed under OHSU IRB study #3330. Patient clinical information and tissues were obtained with informed consent in accordance with the Declaration of Helsinki.

### sci-ATAC-seq specimen processing and library generation

Patient tissue samples were obtained as frozen enzymatic disaggregates except for one frozen whole-viable sample. All samples were thawed on ice and each washed with 1x PBS with Protease inhibitor and centrifuged. The supernatant of the whole-viable sample was removed and treated with 50 uL of Collagenase I, 50 uL of Collagenase, and 240 uL of Hyaluronidase. For all samples, 950 uL of 1x PBS was added, and samples were washed and spun down again. 2 mL of Nuclear Isolation Buffer (NIB) was added on ice for 15 minutes. Samples were then Dounce homogenized with a B pestle on ice. After this, sci-ATAC-seq protocol was performed as previously described [68].

### scRNA-seq specimen processing and library generation

#### Patient tumor tissue enzymatic digestion

Single-cell suspensions from patient tumor tissue were obtained by enzymatic digestion. Briefly, tissue was finely minced using a sterile scalpel for 5 minutes in continuously regenerating cell (CRC) media, followed by a 2-4-hours of enzymatic digestion with Liberase TH (10 µg/ml) (Roche) and ROCK inhibitor (10 µM) (MedChemExpress) at 37 °C using continuous stirring conditions until no visible tissue was present. Two samples went through the same protocol but did not receive ROCK inhibitor: ST-00018963-T and ST-00018724-T. Single cell suspensions were collected by centrifugation at 1000 rpm for 5 minutes and washed with Human Wash Medium. Cells were counted and viability assessed using a Countess 3 (Invitrogen). Cells were then frozen in 90% FBS 10% DMSO and stored at-80 °C.

#### Single-cell RNA sequencing library construction

Frozen single-cell suspensions were thawed at 37 °C and resuspended in DMEM with 2% FBS and ROCK inhibitor. Cells were spun down at 1000 rpm for 5 min at 4 °C and resuspended in PBS with 2% FBS and ROCK inhibitor, then assessed for cell number and viability. About 34,000 cells were loaded onto the Chromium Controller (10x Genomics) for an estimated recovery of 20,000 cells. Library preparation was done with the Chromium Single Cell 3’ v3.1 (10x Genomics) following the manufacturer’s protocol. Initial resultant cDNA was profiled on a 2100 Bioanalyzer (Agilent) before continuing with library preparation. Final libraries were then assessed on a 4200 TapeStation (Agilent) quantified by real-time PCR using the Kapa Bioscience NGS Library Quantification kit on a StepOnePlus Real Time PCR Workstation (Thermo/ABI) and sequenced on a NovaSeq 6000 (Illumina). All FASTQ files were prepared from the raw base call files using bcl2fastq (Illumina). Single cell RNA sequencing assays were performed by the OHSU Massively Parallel Sequencing Shared Resource.

#### CRC media

70% DMEM, 25% F-12, 5% FBS, 4 µg/ml Hydrocortisone, 5 µg/ml Insulin, 8.4 ng/ml Cholera toxin, 10 ng/ml EGF, 24 µg/ml Adenine, 1x Primocin, 10 µM ROCK inhibitor.

#### Human Wash Medium

Advanced DMEM/F-12, 10mM HEPES, 1x GlutaMAX, 100 µg/ml Primocin, 0.1% BSA.

### Sequencing read processing and alignment

#### sci-ATAC data processing and alignment

Reads for all sci-ATAC-seq sequencing runs were demultiplexed and matched to a whitelist of possible barcodes via unidex (https://github.com/adeylab/unidex). Unidex was run under default configurations for single-cell combinatorial indexing (-M sci) and was permitted two Hamming edit distances for barcode matching. Resulting FASTQ files were aligned to the hg38 analysis set using the scitools (https://github.com/adeylab/scitools) wrapper function “fastq-align”, which calls bwa mem [69] (v0.7.17-r1198-dirty) to perform alignment. Aligned BAM files for all sequencing runs were name-sorted, merged, and filtered for q30 quality reads via samtools (v1.16.1) [70]. Finally, PCR-duplicated reads were purged from the unified BAM file in a barcode-aware manner through the scitools function “bam-rmdup”.

#### scRNA-seq data processing and alignment

FASTQ reads were aligned to the GRCh38 reference and subsequently quantified into a counts matrix by Cellranger (v7.0.0) (10x Genomics) through the function “cellranger count”. Intronic reads were included.

### Single cell quality filtering and doublet removal

#### sci-ATAC-seq doublet removal

Aligned sci-ATAC-seq data were read and analyzed by the ArchR R package [71]. We configured ArchR to retain only cell barcodes that had least 1k fragments and 2.5 TSS enrichment. Tumor samples were split into separate ArchR projects, grouped according to which samples were assayed in-multiplex with each other during sci-ATAC-seq preparation. Doublets were identified in each ArchR project/multiplex group via “addDoubletScores” and removed. If fewer than 10% doublets were estimated, the filter ratio (“filterDoublets”) was elevated. All ArchR projects were then re-integrated into a single project containing all passing cells from all samples.

#### sci-ATAC-seq per-cell quality filters

The ArchR project containing all cells from all samples (minimum 1k fragments, 2.5 TSS enrichment) was analyzed at a precursor level before applying additional quality filters. We reduced data dimensionality via ArchR’s iterative LSI method (“addIterativeLSI”; iterations=3, resolution=2, varFeatures=100000) and identified clusters through the ArchR function “addClusters”. Cells that did not have at least 40% of their read content overlapping with reference atlas DNAse hypersensitive sites (DHSs) [72] were removed. This metric is similar to a common ATAC-seq and ChIP-seq signal-to-noise ratio, the fraction of reads in peaks (FRIP), only here we use *a priori* knowledge of known regulatory sites to make this calculation. Passing cells were then re-analyzed using the same addIterativeLSI and addClusters settings. After inspecting this model, we removed stromal and immune cells with less than 4 TSS enrichment, and performed iterative LSI and clustering again. For all focused analyses of primary tumor stromal or immune cells in this work, we created a separate ArchR project from which all metastatic samples were absent; we then performed the same iterative clustering and per-cell filtering workflow described above with the same quality cutoffs.

#### scRNA-seq quality filters, doublet removal, and integration

Unfiltered counts matrices generated by Cellranger were analyzed with the R package Seurat v4 [70]. We removed cells that had fewer than 500 unique reads, fewer than 300 unique genes, and greater than 20% mitochondrial reads. Filtered data were then exported as a counts matrix and doublets were identified from this using the python package Scrublet [73] using an expected doublet rate of 16% and otherwise default parameters. Data were removed of doublets and imported back to R as a Seurat object. We applied the Seurat “SCTransform” v2 workflow using the first 30 principal components and clustered single cells using a resolution of 0.8. Samples were integrated using the SCT integration workflow, briefly: selectIntegrationFeatures using the top 3000 variable features, PrepSCTIntegration, FindIntegrationAnchors (normalization set to “SCT”, and k.anchor set to 5, and reduction set to “rpca”), and finally IntegrateData. We ran “RunPCA”, “FindNeighbors” with dims=1:30, and “FindClusters” on the integrated assay of the Seurat object. Clusters were cell typed using canonical gene expression markers. For sub-clustering of stromal cells, we subset the integrated Seurat R object to contain only stromal cells and re-ran “runPCA”, “FindNeighbors”, and “FindClusters”. After removing two putative doublet clusters with cross-expression of immune and epithelial markers, we re-ran “VariableFeatures”, “RunPCA”, “FindNeighbors”, and “FindClusters” using the integrated assay of the Seurat object, then normalized and scaled the RNA assay of the Seurat object.

### Peak calling

For all analyses focused on epithelial and malignant cells in our sci-ATAC-seq data, we used a background peak set identified by MACS3 [74]. We applied MACS3 to a filtered BAM file containing all primary tumor and metastatic cells (of any cell type) that passed quality control (TSS enrichment > 2.5 in epithelial/malignant, TSS enrichment > 4 in stromal/immune, fragments > 1k, and > 40% reference DHS overlap). MACS3 was set with the following parameters:-f BAMPE-g hs --keep-dup all --min-length 100. The MACS3 peak set contains n=325,149 peaks of variable peak width.

For analyses focused on primary tumor stromal or immune cells, we used a background peak set identified by the ArchR implementation of MACS2. We ran the ArchR functions “addGroupCoverages” and “addReproduciblePeakSet” with “groupBy=’Clusters’“, specifically using the ArchR project containing only primary tumor cells (of any cell type) with sufficient quality metrics. The MACS2 peak set contains n=267,129 peaks of fixed width (500 bp).

### *Cis*-regulatory topic modeling and analysis

#### Topic modeling

To perform single-cell ATAC-seq topic modeling, we applied the R implementation of cisTopic [24] on 1) primary and metastatic epithelial/malignant cells, 2) primary tumor stromal cells, and 3) primary tumor immune cells. For cells in 1), we generated an ATAC-seq peaks counts matrix from the MACS3 peak set (n=325,149 peaks) and used this as input into cisTopic. For 2) and 3), we generated separate counts matrices representing stromal cells and immune cells using the MACS2 peak set (n=267,129 peaks). We then ran the cisTopic workflow on each counts matrix separately. First, we ran “runWarpLDAModels” under default alpha and beta parameterization to identify the optimal topic number. For the stromal and immune topic models, we tested topic numbers between 10 and 50. For stromal cells, we selected the 27-topic model based on its high second derivative on the log-likelihood curve (Extended Fig. 8a). For immune cells, we initially attempted to use a high second-derivative 16-topic model, but analysis of this model did not achieve adequate resolution of immune cell states. We therefore selected the model with the lowest perplexity score, a 43-topic model (Extended Fig. 9a). For the epithelial/malignant cisTopic model, we tested a larger range of topic numbers (between 10 and 100) to accommodate potential patient-specific malignant states. We ultimately selected the 50-topic model due to its low perplexity (Supplementary Fig. 2a) and high resolution of cell states. Finally, to define which chromatin regions belong to individual topics, we ran “getRegionScores” with method=”NormTop” and “binarizecisTopics” with thrP=0.975 on each cisTopic model.

#### Single cell clustering and UMAP embedding

We also used cisTopic models to perform single cell clustering and UMAP embedding of epithelial/malignant cells (Fig. 1e), stromal cells (Fig. 5a), and immune cells (Extended Fig. 9b). Per-cell scaled topic score matrices were exported used as input for the R packages umap (v0.2.10.0) and PhenoGraph (Rphenograph; v0.99.1) [75]. PhenoGraph clustering *k* was set to 50 for the immune topic model, and to 15 for the stromal and epithelial/malignant topic models.

#### Identification of malignant subtype-associated topics

To identify regulatory topics that are enriched in SIP, classical-NRP, or basal committed cells across primary PDAC tumors (excluding PAV tumors), we took a two-step analytical approach:

First, we collapsed topic scores from per-cell values to sample-level means. This prevents single cells from counting as individual samples (*n*) in statistical testing, which can lead to overestimation of significance. To collapse our data to sample-level while also preserving intra-tumor heterogeneity, we split the malignant population of each sample by subtype (SIP, C-NRP, BC), and calculated mean topic scores within each. Within each sample, we excluded any subtype subpopulation with fewer than 10 cells. We retained only subpopulations from primary PDAC tumors and excluded PAV and metastatic samples.

Second, we modeled the mean topic scores through a linear mixed-effects framework. We modeled each topic individually, treating malignant subtype as a fixed effect and tumor sample as a random effect. The random intercept accounts for the shared environment and processing of subtypes originating from the same tumor (intra-sample correlation). For each topic, we applied the R package lme4 (v1.1-29) and modeled mean topic scores using the formula: *(mean topic scores) ∼ subtype + (1 | sample)*. We then calculated estimated marginal means via the R package emmeans (v1.11.1) and conducted pairwise comparisons between SIP, basal committed, and classical-NRP (FDR corrected). The topic analyzed in this workflow was deemed subtype-associated if one of the three subtypes had significantly greater marginal means (FDR < 0.05) than both of its counterparts. This workflow initially identified 21 subtype-associated topics, but we excluded two from further analysis: Topic 19, which was specific to one sample (ST-00004898-M); and Topic 24, which was specific to pulmonary epithelial cells (Extended Fig. 3a).

#### Enrichment of published PDAC and CAF gene signatures within topics

cisTopic provides a GSEA-like algorithm (“getSignaturesRegions”) to score enrichment of epigenomic signatures within topics. To analyze topic enrichment of published gene signatures, we converted the gene signatures into proxy peak sets suitable as input for getSignaturesRegions. Proxy peak sets for each PDAC subtype signature were generated as follows: first, we paired all MACS3-called peaks (n=325,149 peaks) to their nearest gene TSS via ChIPSeeker [76]. Second, for each subtype gene signature, we created a peak set containing any peak that was paired to a gene in the original signature. Third, we refined the peak sets by excluding any peak that was more than 50 kb away from its paired gene. For CAF signatures, we applied the same workflow, but used the MACS2-called peak set (n=267,129 peaks) to generate proxy peak signatures.

#### Epithelial cell enrichment of neural-like progenitor and neuroendocrine signatures

Per-cell enrichment of proxy peak sets for NRP and NE PDAC signatures [7] was performed using the cisTopic implementation of AUCell [77]. Cell ranking and signature enrichment was performed via the functions “AUCell_buildRankings” and “signatureCellEnrichment”.

### Differential accessibility testing

All differential accessibility (DA) tests throughout this work were run in ArchR using the function “getMarkerFeatures”. We applied the binomial test function of getMarkerFeatures using the default settings for this test method. All DA peaks identified from our PDAC sci-ATAC-seq dataset are defined as |log2 fold change| > 1.0 and FDR < 0.01. For the published pancreatic stellate cell *in vitro* dataset by Bowman CL and Daniel CJ, et al. [61], DA peaks are defined as |log2 fold change| > 1.0 and FDR < 0.05.

### TF motif enrichment

For per-cell TF motif accessibility scores, we applied the ArchR implementation of chromVAR, which Z-scores the deviation of each cell’s TF motif accessibility from the global average [78]. For enrichment of TF motifs in epithelial/malignant topics, we applied the ArchR function “peakAnnoEnrichment” to count the number of peaks that contain each TF motif. We compared motif counts in topic peaks to background (all unique peaks across all 50 topics in the cisTopic model) by hypergeometric test using the R function “phyper”. Log10(*p*) values from all tests across all 50 topics were Bonferroni-corrected. TF motif analyses in single cells and in epithelial/malignant topics both utilized the “cisbp” motif set provided in the ArchR implementation of chromVAR.

To identify TF motifs enriched in stromal topics, we applied the HOMER motif discovery algorithm [79]. We ran the HOMER wrapper program “findMotifsGenome” (http://homer.ucsd.edu/homer/ngs/peakMotifs.html) to test enrichment of TF motifs from the HOMER “known motif” library. The background for these tests was the collection of all unique peaks across all 27 topics in the stromal cisTopic model. P-values calculated by HOMER for all TF motif tests across all 27 stromal topics were FDR-corrected.

### Estimation of copy number alterations

We applied the R package epiAneufinder (v0.1.0) to estimate copy number changes in our sci-ATAC-seq dataset [80]. To generate the required epiAneufinder input, we converted our sequencing data from BAM format to scATAC-seq fragments format using Sinto (v0.9.0) (https://github.com/timoast/sinto). We then ran the “epiAneufinder” function with windowSize=10000 on all quality-filtered cells from all samples in our sci-ATAC-seq fragments file. We excluded blacklisted regions in the hg38 genome (see https://github.com/colomemaria/epiAneufinder).

### Annotation of malignant epigenetic subtypes

#### Cluster annotation

We classified malignant epithelial clusters as *GATA6^-^*if they showed significantly lower *GATA6* accessibility than the acinar/ductal cluster (Wilcoxon FDR < 0.05, log2 fold change of mean gene score <-0.5). Clusters that did not meet this criterion were deemed *GATA6^+^*. Likewise, we classified malignant clusters as *KRT17^+^* if they showed significantly greater *KRT17* accessibility than the acinar/ductal cluster (Wilcoxon FDR < 0.05, log2 fold change of mean gene score > 0.5); clusters that did not meet this criterion were deemed *KRT17^-^*. The *GATA6* and *KRT17* gene accessibility scores used in these tests were calculated by ArchR (“addGeneScoreMatrix”) and represent per-cell depth-normalized quantitative approximations of the overall accessibility within and around genetic loci. *KRT17^+^ GATA6^-^* malignant clusters were annotated as basal committed, *GATA6^+^ KRT17^-^* malignant clusters were annotated as classical-NRP, and *GATA6^+^ KRT17^+^* co-accessible malignant clusters were annotated as subtype-intermediate progenitor.

#### Sample annotation

Each tumor in our sci-ATAC-seq cohort was classified as SIP, classical-NRP, or basal committed subtype according to the tumor’s most numerically prevalent malignant subpopulation. Two primary PDAC samples were tied between SIP and another subtype by cell count: ST-00017762-T and ST-00020291-T. In both cases, we selected the non-SIP subtype as the representative subtype of the sample, and deemed ST-00017762-T as basal committed and ST-00020291-T as classical-NRP. We did not assign a tumor epigenetic subtype to samples with fewer than 10 malignant cells.

### Lineage-specific enhancer enrichment

Reference regulatory regions for human cell types and lineages were obtained from the sci-ATAC-seq atlas published by Zhang K and Hocker JD, et al. [25]. This dataset provides 150 *cis*-regulatory “modules”, which are clustered sets of regulatory regions enriched in certain lineages (e.g. pancreatic epithelial, gastric epithelial). It also provides accessible regions identified in 222 cell types.

#### Epithelial cell typing

To identify epithelial cell types, we analyzed per-cell accessibility of epigenomic signatures from the “module” collection by Zhang K and Hocker JD, et al. [25]. Per-cell accessibility scores for these modules were calculated via the ArchR implementation of chromVAR. We selected 5 representative modules for data visualization of cell types: module #72 “pancreatic epithelial”, #72 “gastrointestinal epithelial”, #85 “neuroendocrine”, #93 “gastric epithelial”, and #65 “pulmonary epithelial”. Following cell type annotation, cells from primary tumors that co-clustered with pulmonary epithelial cells were excluded (n=21)). Likewise, cells from metastases that co-clustered with non-malignant epithelial cell types (except pulmonary epithelial) were excluded (n=83).

#### Cell type enrichment within topics

For analysis of lineage enhancers in subtype-associated topics, we utilized the 222 cell type peak sets by Zhang K and Hocker JD, et al. [25]. We counted the overlap of cell type peaks with 1) peaks in individual topics, and 2) the background set of all topic peaks in the cisTopic model. For each topic, we performed hypergeometric tests for the 222 cell types using the R function “phyper” and applied Bonferroni correction to log10(*p*) values from all tests across all 50 topics.

### Gene Ontology enrichment in topics

Enrichment of GO biological process terms in topics was determined via GREAT [81]. Peaks from an individual topic were used as the test set. The background set was all unique topic peaks from the relevant cisTopic model (epithelial/malignant, stromal, or immune).

### Module scoring of PDAC and CAF subtype signatures

To quantify the per-cell epigenetic module scores for basal-like, classical, mesenchymal PDAC gene signatures as well as CAF gene signatures, we applied the ArchR function “addModuleScore” with default settings. This function takes a list of genes and combines their individual signals (equally weighted) into a single quantitative “module” score (https://www.archrproject.com/reference/addModuleScore.html). Per-cell transcriptomic module scores were calculated using the Seurat function “AddModuleScore” using SCT gene expression values.

### Correlation between immune topics and malignant topics

To correlate malignant subtype-associated topics with immune topics, we first collapsed topic scores into sample-level means. For each primary PDAC sample, we calculated mean immune topic scores within every immune cell type present with at least 10 cells in the sample. For each primary PDAC sample with at least 10 malignant cells, we also calculated the mean scores of subtype-associated topics from all malignant cells in the sample. We then calculated Kendall correlation coefficients using the mean immune topic scores and mean malignant subtype topic scores of each sample. Samples that did not have sufficient cell counts for both populations (immune cell type and malignant cells) or correspond to PAV tumors were excluded.

### Cyclic immunofluorescence analysis

Multiplex immunofluorescence images of SIP, BC, and C-NRP tumor sections of were obtained from our pre-existing tumor microarray, which was previously generated and imaged as described in Link JM, et al. [21]. As cited [21], unsupervised clustering of single-cell mean intensity was used to define cell types, using the Leiden algorithm implemented in scanpy (v.1.9.3) [82]. For marker positivity on each cell type, each autofluorescence subtracted marker was visualized in napari and a threshold was selected to exclude all background pixels. T cells were defined by Leiden clustering, and positivity for the following markers was defined as: mean intensity > 1024 for CD90, > 1536 for PD-1, > 1152 for CD4, and > 3072 for FOXP3. Immune cells were defined by Leiden clustering, and were deemed PD-L1^+^ if mean PD-L1 intensity was > 50.

### Curation of published gene signatures

We curated gene signatures for PDAC subtypes by Bailey P, et al. [38] from the differential expression tables provided in their publication. From their differential expression results that compared each subtype (pancreatic progenitor, squamous, ADEX, and immunogenic) to the rest, we collected the top 200 genes with the highest log fold change value (adjusted p-value < 0.05). All other PDAC subtype gene signatures were collected as provided by their original publication [2, 3, 18, 29]. For PSC and CAF signatures by Wang Y and Liang Y, et al. [53] and Oh K, et al. [6], we collected all significant (adjusted p-value < 0.05) marker genes for each cell type/CAF subtype with a log fold change value > 0.5.

### Development of gene signatures from sci-ATAC-seq data

#### Malignant epigenetic subtypes

We developed gene signatures for epigenetic subtypes from subtype-specific DA peaks. First, we performed pairwise DA tests between SIP, classical-NRP, and basal committed cells from primary PDAC tumors. Second, we intersected the positive DA peak hits for each subtype; peaks that were significantly more accessible (log2 fold change > 1.0, FDR < 0.01) in both comparisons were deemed subtype-specific. This produced peak sets for SIP (n=127 specific peaks), classical-NRP (n=1,265), and basal committed (n=11,896). Third, we linked peaks to their nearest genomic feature via ChIPSeeker [76]. We removed any genomic feature that was more than 50 kb apart from its linked peak, and retained only the features that have an associated Ensembl IDs and are annotated as protein coding, microRNA, or long non-coding RNAs (to exclude unwanted features such as pseudogenes). This produced a SIP signature of 84 genes. However, classical-NRP and basal committed gene sets required trimming due to their excessive size. To rank genes based on importance, we used the predictive distribution of our epithelial/malignant cisTopic model, calculates the likelihood of each peak occurring in cells. We ranked peaks based on the mean probability of a peak occurring in classical-NRP/basal committed cells, then selected the top unique 200 genes based on this ranking. This size is comparable to MSigDB “Hallmark” gene sets [83, 84] and PDAC malignant lineage gene sets by Hwang WL, et al. [7].

### CAF topic signatures

We extracted peaks from Topic 11 (representing iCAFs), Topic 15 (*LRRC15^+^*myCAFs), and Topic 22 (*NGFR^+^* myCAFs). Next, we connected them to their nearest genetic feature via ChIPSeeker [76], discarding any peak-gene linkage that was further than 50 kb from the gene TSS or had no associated Ensembl ID. Using the topic-peak predictive distribution in our stromal cisTopic model, we ranked peaks from most-to-least likely to be associated with the corresponding topic, and from this ranking selected the top 200 associated genes.

### Analysis of TGFβ-responsive PSC enhancers

To identify PSC enhancers that open or close in response to TGFβ, we analyzed previous sci-ATAC-seq data by Bowman CL and Daniel CJ, et al. [61]. We collated the post-filtered sci-ATAC sequence data and called peaks from this publication into an ArchRProject for ArchR analysis. We examined differential accessibility between the two short hairpin scramble (shSCR) control groups from this publication (TGFβ-treated shSCR PSCs vs. untreated shSCR PSCs) and excluded experimental groups with PIN1 knockdown (TGFβ-treated shPIN1 PSCs and untreated shPIN1 PSCs). Differentially accessible peaks from these data (FDR<0.05 and log2 fold change>1.0) were identified using the binomial test settings for ArchR “getMarkerFeatures”. In our human PDAC sci-ATAC-seq dataset, per-cell accessibility of the PSCs peaks was scored using the ArchR implementation of chromVAR.

### Gene signature scoring in bulk RNA-seq data

We analyzed bulk RNA-seq profiles for our previous cohort of 218 human primary pancreatic tumors, described previously [21]. To do so, we collated transcript abundance outputs for all primary tumors via tximport (v1.22.0) [85] and then created a gene-level object for DESeq2 (v1.34.0) [86] via *“*DESeqDataSetFromTximport” Genes with fewer than 10 cumulative reads across all 218 samples in the dataset were removed. For gene set analysis, we applied variance-stabilizing transformation (VST) to expression counts using DESeq2. VST counts were then used to perform gene set variation analysis (GSVA) using the GSVA R package [87].

### Survival analysis

Patient survival information (days from diagnosis to follow-up; vital status at follow-up) were obtained from our previous publication [21]. To calculate univariate hazard ratios for gene expression signatures, we used GSVA scores to model survival using the “surv” and “coxph” functions of the R package survival (v3.3-1). To compare the survival curve of SIP tumors to other epigenetic subtypes in our sci-ATAC-seq cohort, we used the survminer (v0.4.9) R package. Survival curves were visualized via “ggsurvplot” and log-rank tests were performed via “pairwise_survdiff”.

### Data visualization and plotting

All heatmaps presented in this work were created with the R package pheatmap (v1.0.12), with the exception of the stromal marker peak heatmap which was generated via the ArchR function “plotMarkerHeatmap”.

Continuous color scales used this work are applied from color palettes in the R packages ArchR, RColorBrewer, and Seurat. Genomic track views/UCSC-style genome browser plots were generated using the ArchR function “plotBrowserTrack”. UMAPs were plotted using the ArchR function “plotEmbedding” and adjusted using ggplot2 (v3.4.1). All other plots were generated using ggplot2. Data denoising and imputation were performed using the ArchR implementation of MAGIC (Markov Affinity-based Graph Imputation of Cells) [88] using iterative LSI matrices (for ArchR sci-ATAC-seq models), or scaled cisTopic matrices (for cisTopic sci-ATAC-seq models).

## Data Availability

Raw DNA/RNA sequence data for sci-ATAC-seq and scRNA-seq profiles will be deposited in the controlled access database dbGaP through the NCI Genomic Data Commons. Fragment files for sci-ATAC-seq profiles and gene counts matrices for scRNA-seq profiles (which both provide analyzable data without sequence information) will be deposited in the GEO database. Processed sci-ATAC-seq data for epithelial, stromal, and immune cells will be uploaded to Zenodo as “ArchRProject” folders, which can be imported into R with the ArchR package and subsequently analyzed. Sci-ATAC-seq topic models for epithelial, stromal, and immune cells will also be uploaded to Zenodo as cisTopic R objects.

## Code Availability

Code used to produce analyses and figures in this manuscript will be made available at https://github.com/k-hawth/PDAC_sciATAC.

---

**Extended Figure 1:**
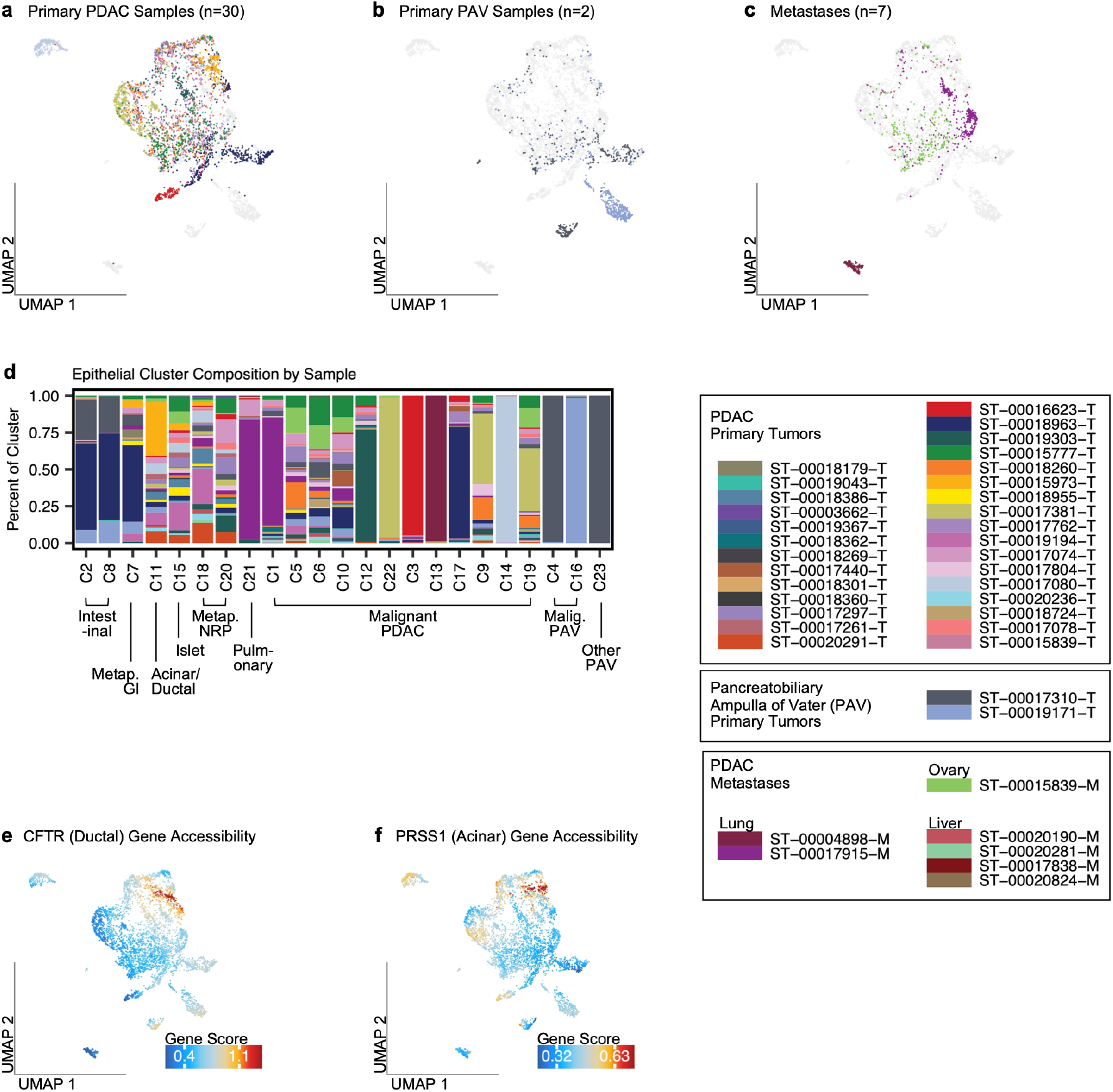
Sample contribution to epithelial/malignant clusters. **a)** UMAP of epithelial/malignant cells. Cells depicted in color belong to a primary PDAC tumor; cells depicted in gray belong to other samples (primary PAV or metastases). **b-c)** Same as panel A, but for primary PAV samples and metastases. **d)** Sample composition of epithelial/malignant clusters. **e-f)** Imputed chromatin accessibility scores for the ductal marker *CFTR* and the acinar marker *PRSS1* visualized on UMAP. Colors are scaled from the 0th to 99th score percentile.

**Extended Figure 2:**
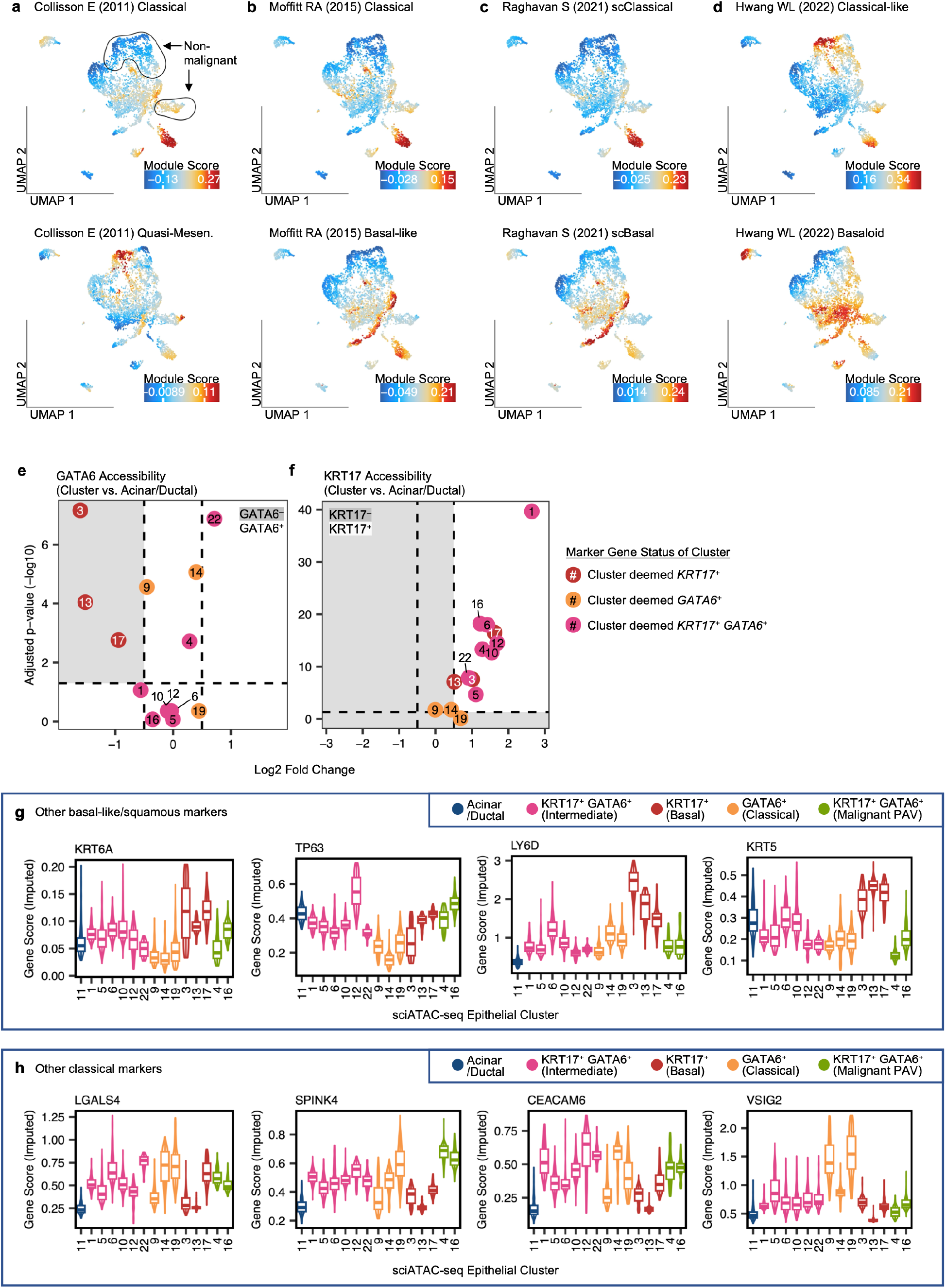
Annotation of malignant clusters as *GATA6^+^* or *KRT17^+^*. **a-d)** Epithelial/malignant UMAP depicting imputed per-cell module scores for canonical basal-like and classical PDAC signatures. **e)** Determination of cluster GATA6 status. GATA6 gene accessibility scores were compared between each malignant cluster and acinar/ductal cells (Wilcoxon test). Malignant clusters are represented as dots, and the number of the cluster is shown within the dot. Clusters that fall within the gray panel (FDR<0.05, mean log2 fold change<-0.5) were deemed GATA6 negative. **f)** Same as panel e), but for KRT17. Clusters that fall within the white panel (FDR<0.05, mean log2 fold change>0.5) were deemed KRT17 positive. **g-h)** Imputed per-cell gene accessibility scores for canonical basal-like and classical markers. Numbers refer to the number of the cluster; colors depict cell type/malignant subtype.

**Extended Figure 3:**
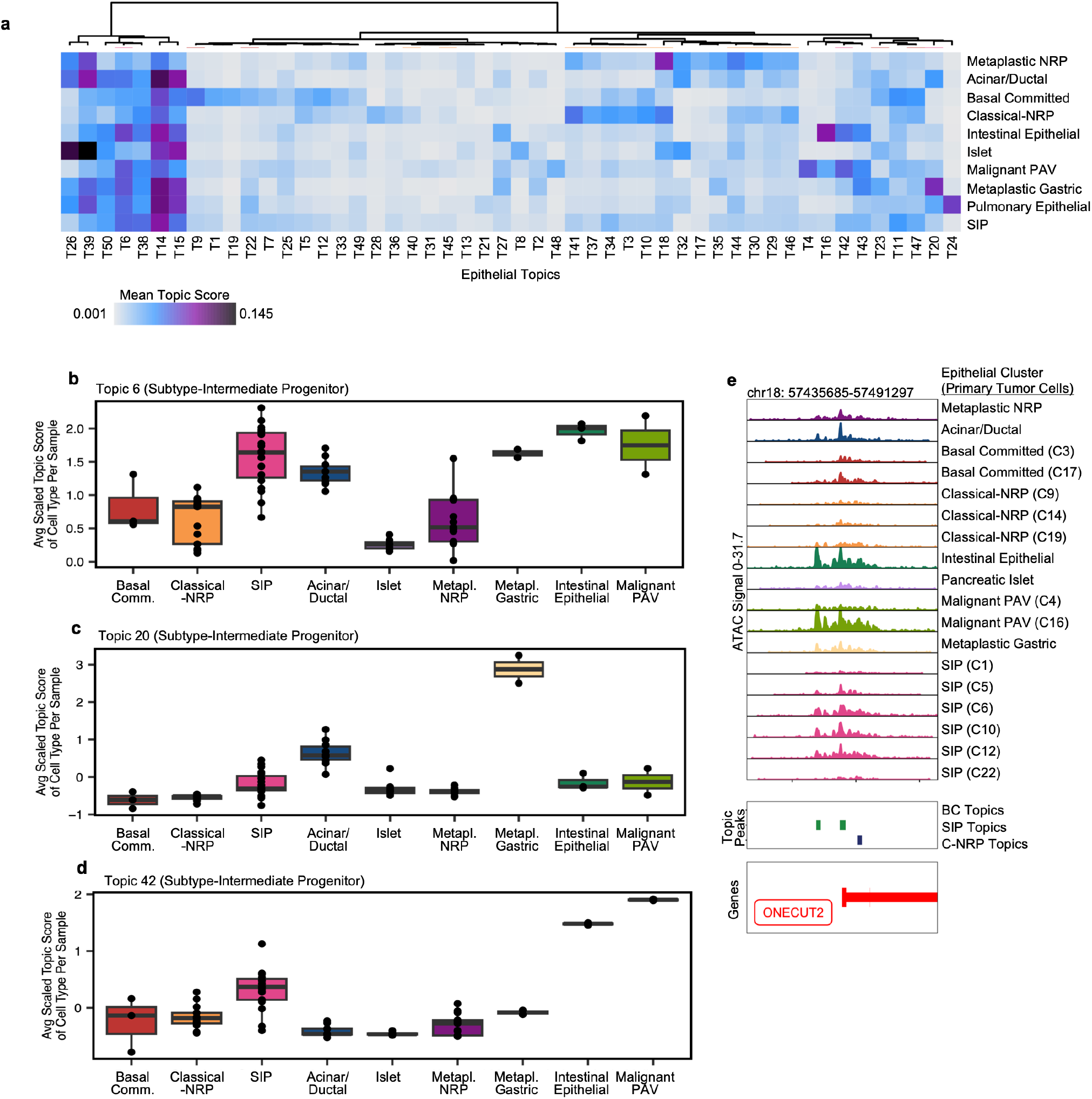
Epithelial/malignant cell type topic scores and SIP-associated topics. **a)** Heatmap of mean topic score by epithelial/malignant state across all samples. **b-d)** Per-sample mean scaled topic scores by cell type. Each point represents the mean topic score of a tumor sample within the cell type depicted on the X-axis. Topic means from all samples shown. **e)** Genomic distribution of ATAC-seq signal near ONECUT2. Rows depict distribution of ATAC-seq signal by cell type. Middle box depicts genomic regions encompassed by basal committed, SIP, and classical-NRP topics.

**Extended Figure 4:**
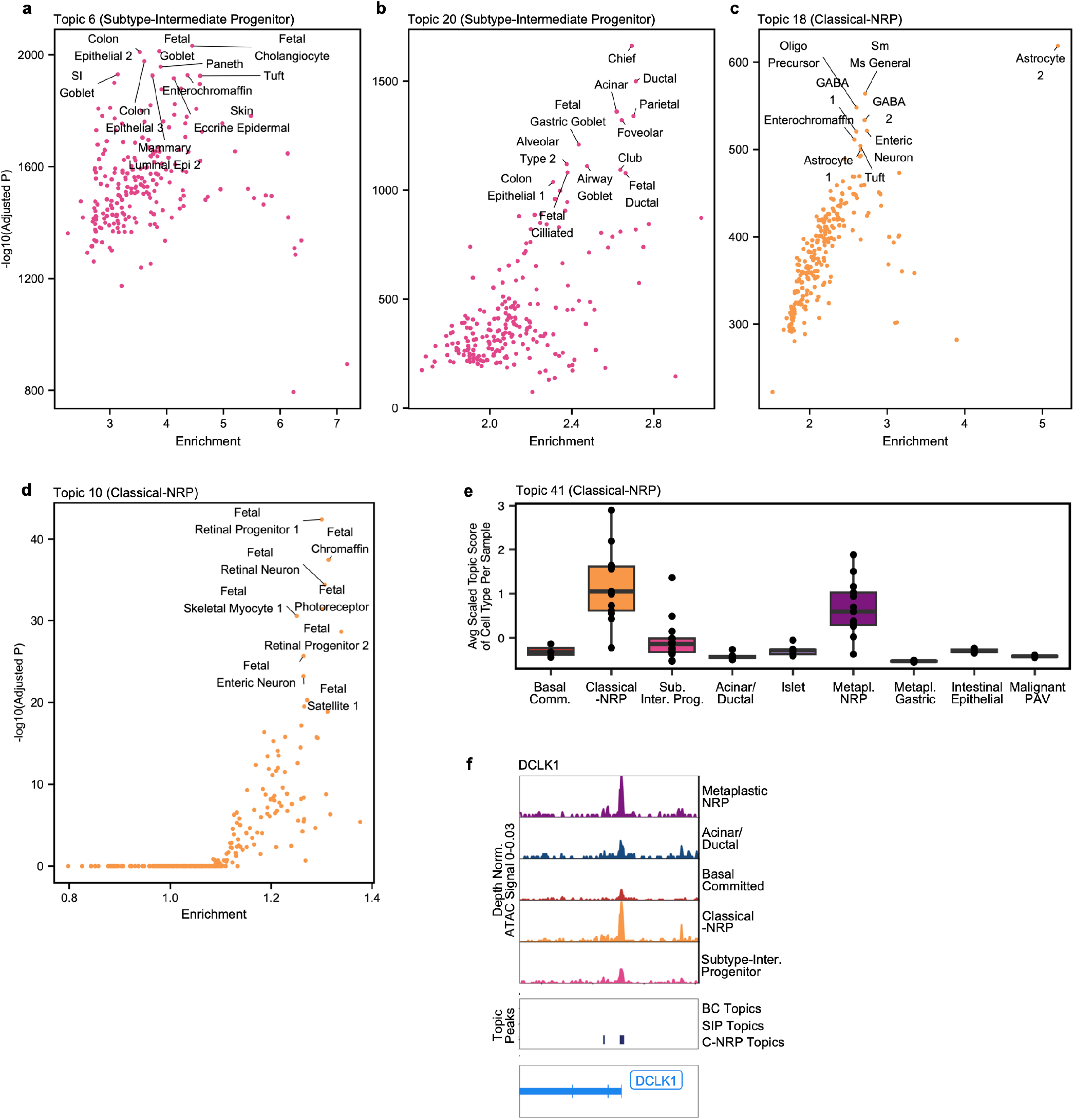
Lineages enriched in other SIP and classical-NRP topics. **a-d)** Per-topic hypergeometric enrichment of cell type enhancers. Each point represents a cell type peak set (222 total) collected from Zhang K and Hocker JD, et al. [25]. Colors refer to the epigenetic subtype associated with the topic. **e)** Per-sample mean scaled topic score for Topic 41 by cell type. Each point represents the mean Topic 41 score of a tumor sample within the cell type depicted on the X-axis. Topic means from all samples shown. **f)** Genomic distribution of ATAC-seq signal near the tuft cell marker *DCLK1.* Rows correspond to cell types. Middle box depicts chromatin regions that are part of subtype-associated regulatory topics.

**Extended Figure 5:**
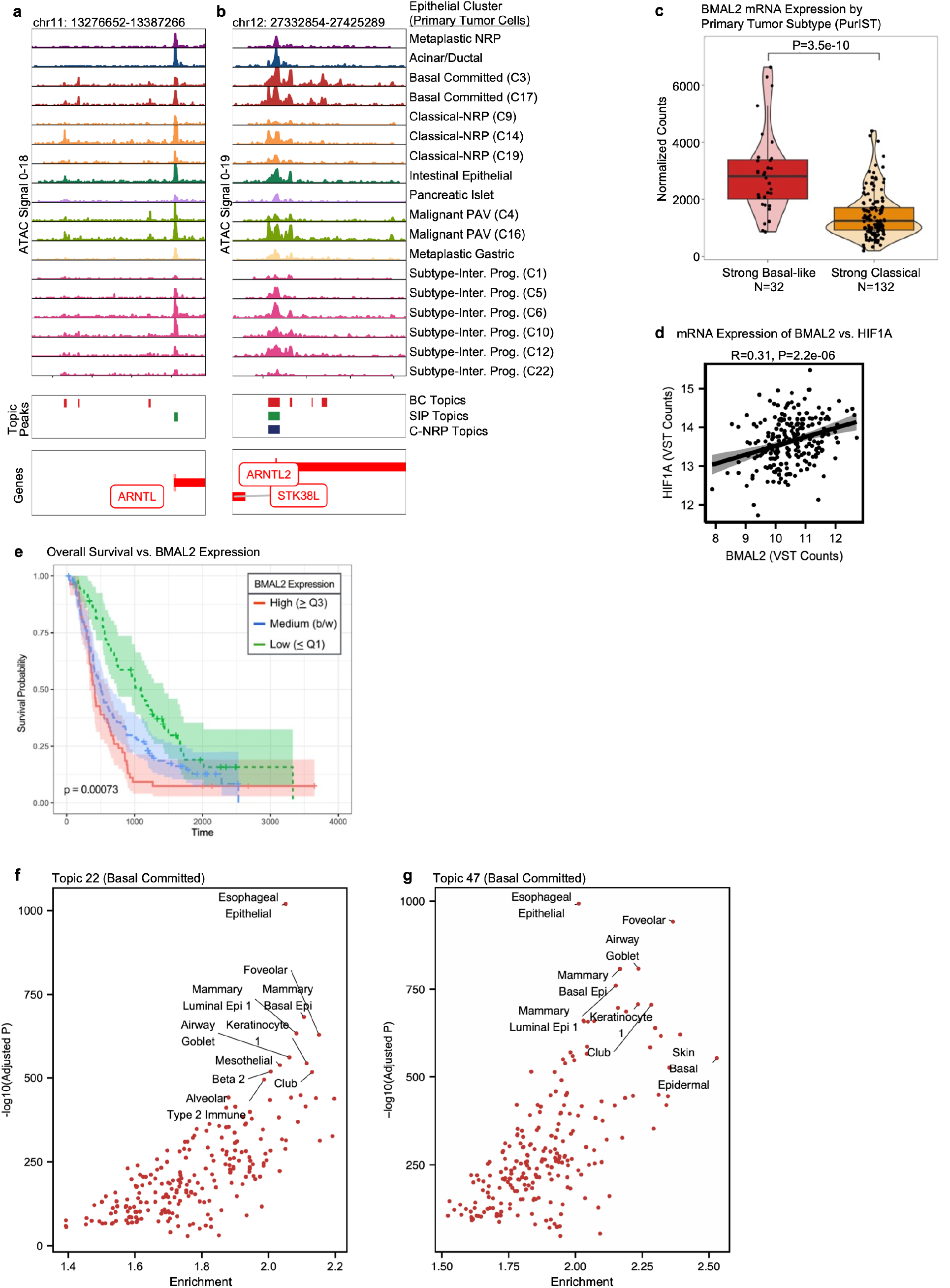
Basal committed subtype is regulated by circadian factors. **a-b)** Genomic distribution of ATAC-seq signal near *ARNTL* and *ARNTL2/BMAL2* in primary tumors. Rows depict distribution of ATAC-seq signal by cell type. Middle box depicts genomic regions encompassed by basal committed, SIP, and classical-NRP topics. **c)** Differential mRNA expression of *BMAL2/ARNTL2* between basal-like and classical primary tumors in the OPTR bulk RNA-seq cohort. **d)** Correlation between *BMAL2/ARNTL2* and *HIF1A* mRNA expression in the OPTR bulk RNA-seq cohort. Primary tumors only. Gene expression scores were transformed with variance-stabilizing transformation (VST). **e)** Overall survival in the primary tumor OPTR cohort by *BMAL2/ARNTL2* mRNA expression. Cohort was divided into the top quartile of *BMAL2* expression (high), bottom quartile (low), and all else (medium). **f-g)** Per-topic hypergeometric enrichment of cell type enhancers in basal committed Topic 22 and Topic 47. Each point represents a cell type peak set (222 total) collected from Zhang K and Hocker JD, et al. [25].

**Extended Figure 6:**
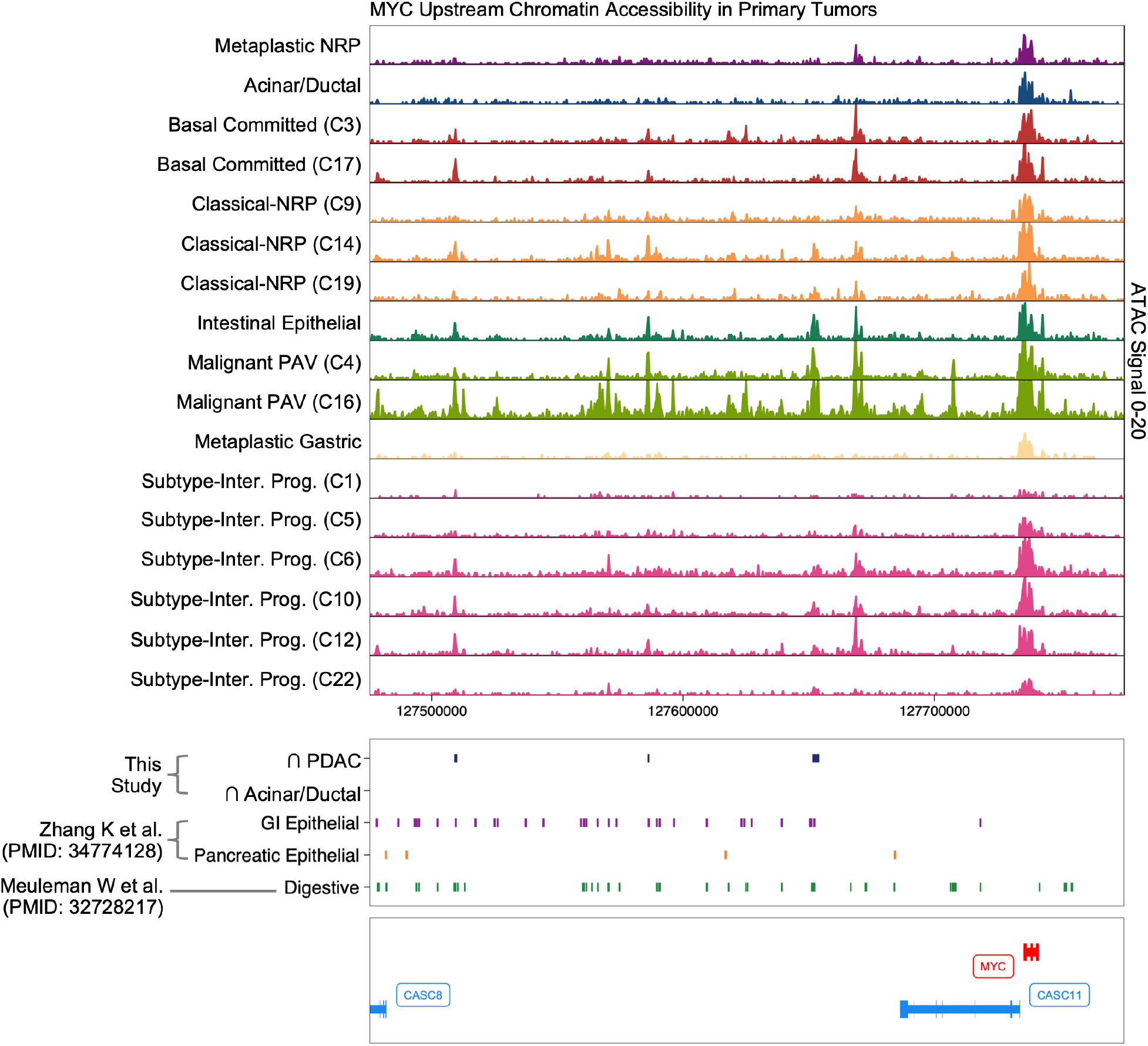
All subtypes activate gastrointestinal enhancers upstream of *MYC*. Genomic distribution of ATAC-seq signal upstream of the proto-oncogene *MYC.* Each row depicts ATAC-seq signal within a specific cluster or cell type. Middle box depicts genomic regions encompassed by consensus PDAC peaks, as well as enhancers identified from normal gastrointestinal epithelial lineages, pancreatic epithelial lineages, and other digestive lineages from reference human atlases.

**Extended Figure 7:**
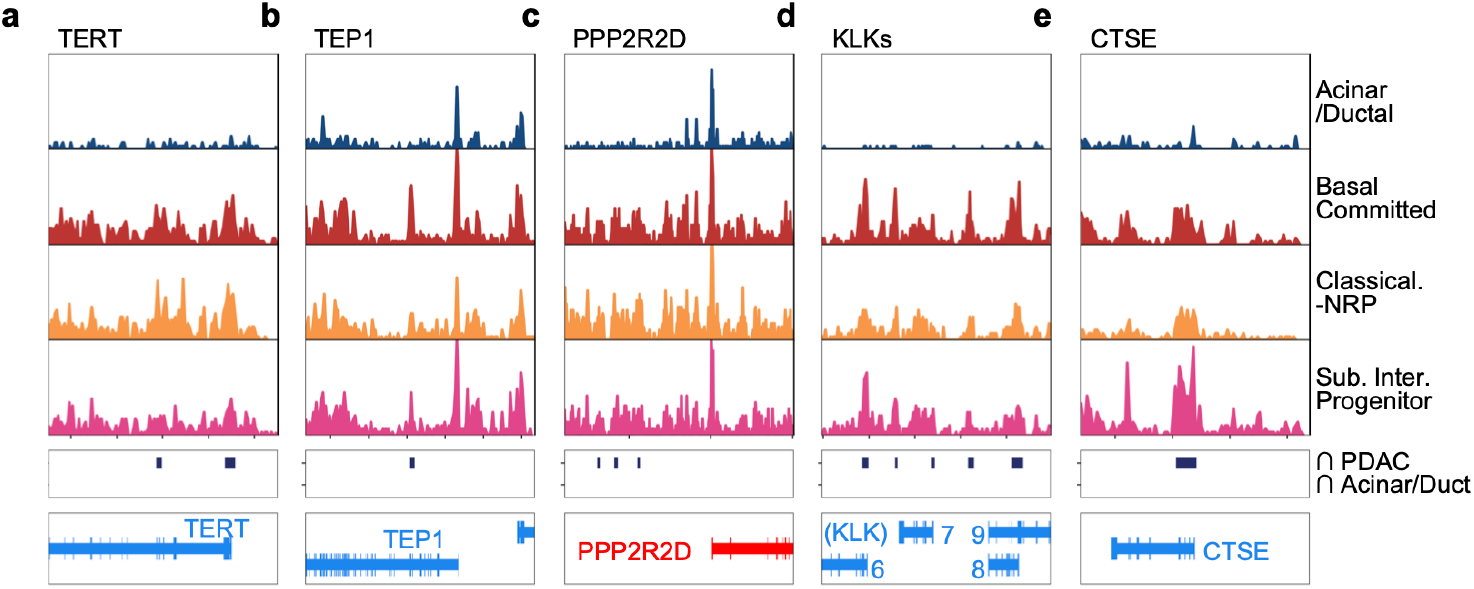
Consensus PDAC peaks near oncogenes. **a-c)** Genomic distribution of ATAC-seq signal upstream of various oncogenes and other factors related to tumorigenesis. Each row depicts ATAC-seq signal within a specific cluster or cell type. Middle box depicts genomic regions encompassed by consensus peak sets (no consensus acinar/ductal peaks exist in the regions shown). Bottom box depicts genomic location of genes. Red genes are encoded on the positive strand; blue on negative. **d)** Same as panels a-c), but depicting the gene cluster of KLKs. The number of each KLK is shown (e.g. *KLK6*, *KLK7*, etc.). **e)** Same as panels a-d).

**Extended Figure 8:**
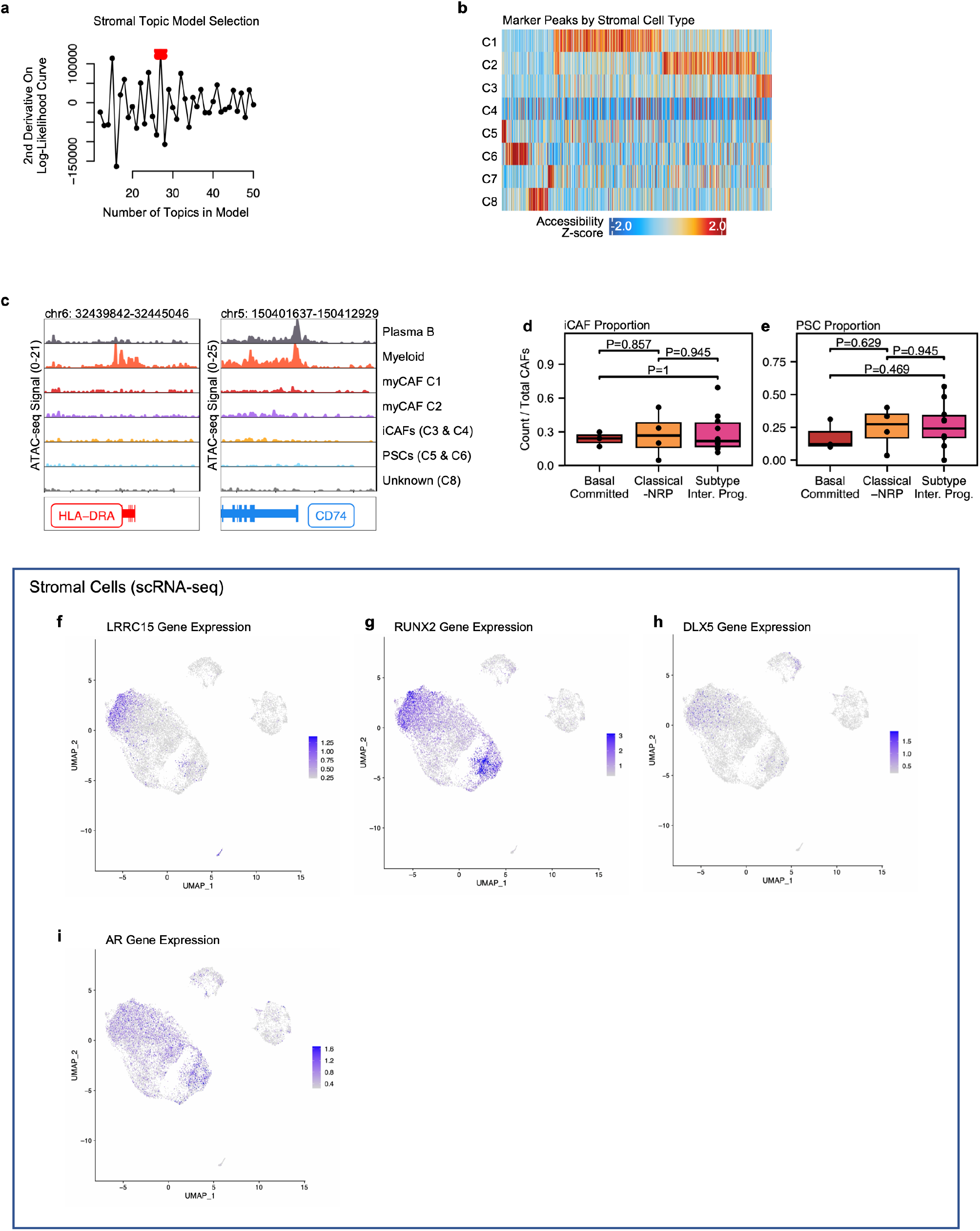
Epigenetic CAF subtype marker gene expression and cell count proportions by malignant PDAC subtype. **a)** cisTopic model selection for primary tumor stromal cells. Red dot indicates the selected model. **b)** Heatmap of significant marker peaks (binomial FDR < 0.05 & log2 fold change > 0.5) identified for each stromal cell type (rows) by ArchR. Columns depict individual peaks. Color scale represents the Z-scored accessibility of the peak. **c)** Genomic distribution of ATAC-seq signal near the MHC-II locus *HLA-DRA* (left) and *CD74* (right). Rows depict ATAC-seq signal from immune cells (Plasma B, Myeloid) and stromal cells (all else). **d-e)** Proportion of iCAFs and PSCs out of the total number of CAFs in primary PDAC tumors. Each point depicts a primary PDAC tumor; the Y-axis depicts the each tumor’s proportion of iCAFs or PSCs in relation to total CAFs. Wilcoxon P-values are shown. **f-i)** Gene expression (log-normalized) of *LRRC15, RUNX2, DLX5,* and *AR* in stromal cell types from scRNA-seq data. Expression values shown are between the 3rd and 97th percentiles.

**Extended Figure 9:**
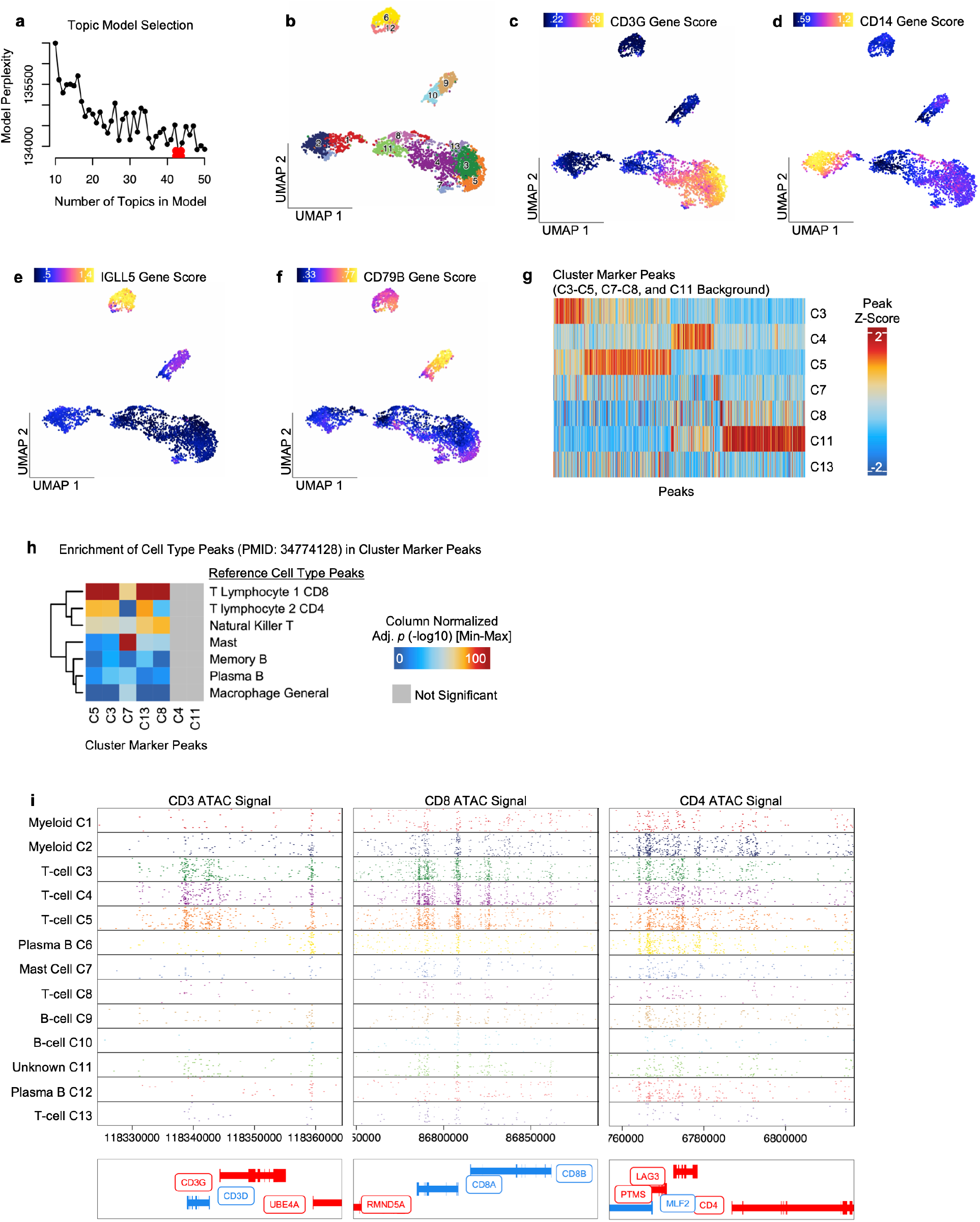
Immune cell typing and topic model selection. **a)** cisTopic model selection for primary tumor immune cells. Red dot indicates the selected model. **b)** UMAP and single cell clustering results for primary tumor immune cells. Colors and numbers refer to individual clusters. **c-f)** UMAP colored by imputed per-cell gene accessibility scores for immune cell type markers. **g)** Heatmap of marker peaks (binomial FDR < 0.05, log2 fold change > 0.5) identified between clusters C3-5, C7-8, and C11. Marker peak analysis performed with ArchR using the function “getMarkerFeatures” and plotted using “markerHeatmap”. Columns depict an individual peak. Color scale represents the Z-scored accessibility of the peak. **h)** Hypergeometric enrichment of reference cell type accessible regions (rows) from [145] in cluster marker peaks from panel g). Color scale values are column-normalized (min-max) Bonferroni-corrected-log10(p-values). **i)** Genomic distribution of ATAC-seq signal near *CD3* genetic loci, *CD8* genetic loci, and the *CD4* locus. Rows indicate ATAC-seq signal from the labeled cluster/cell type. Each point in the plot represents an ATAC-seq read. Bottom box depicts genomic location of genes; red genes are encoded on the plus strand, blue on minus.

**Extended Figure 10:**
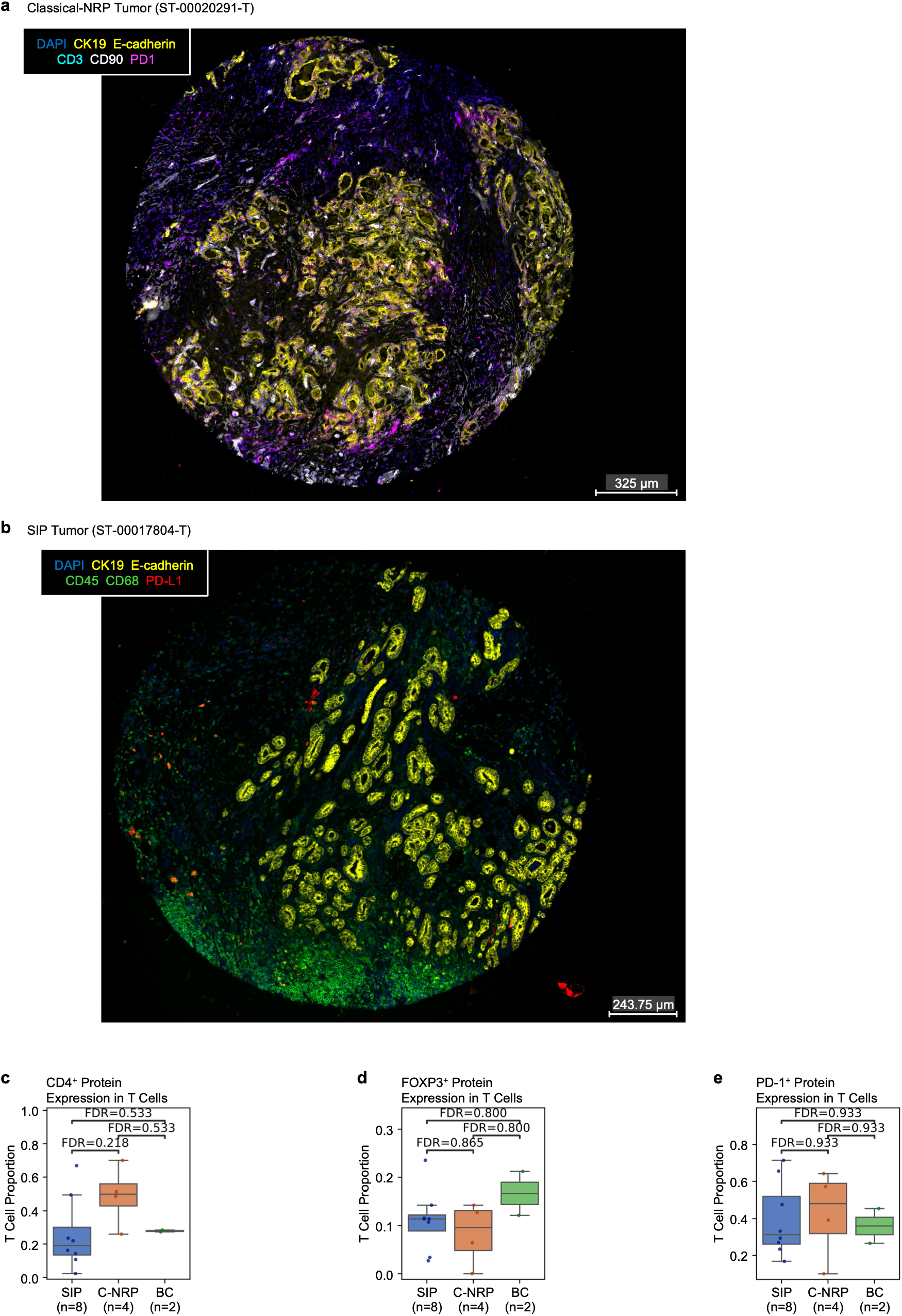
Proportion of CD4^+^, FOXP3^+^, and PD-1^+^ T cells by PDAC epigenetic subtype. **a-b)** Representative classical-NRP and SIP tumor sections, cyclically immunostained for markers depicted at the upper left of each image. **c-e)** Cyclic immunofluorescence analysis of primary PDAC tumor histologic sections. The Y axis depicts the proportion of T cells that show CD4 staining (panel c), FOXP3 staining (panel d), and PD-1 staining (panel e). Significance was determined via Mann-Whitney U test.

**Supplementary Figure 1:**
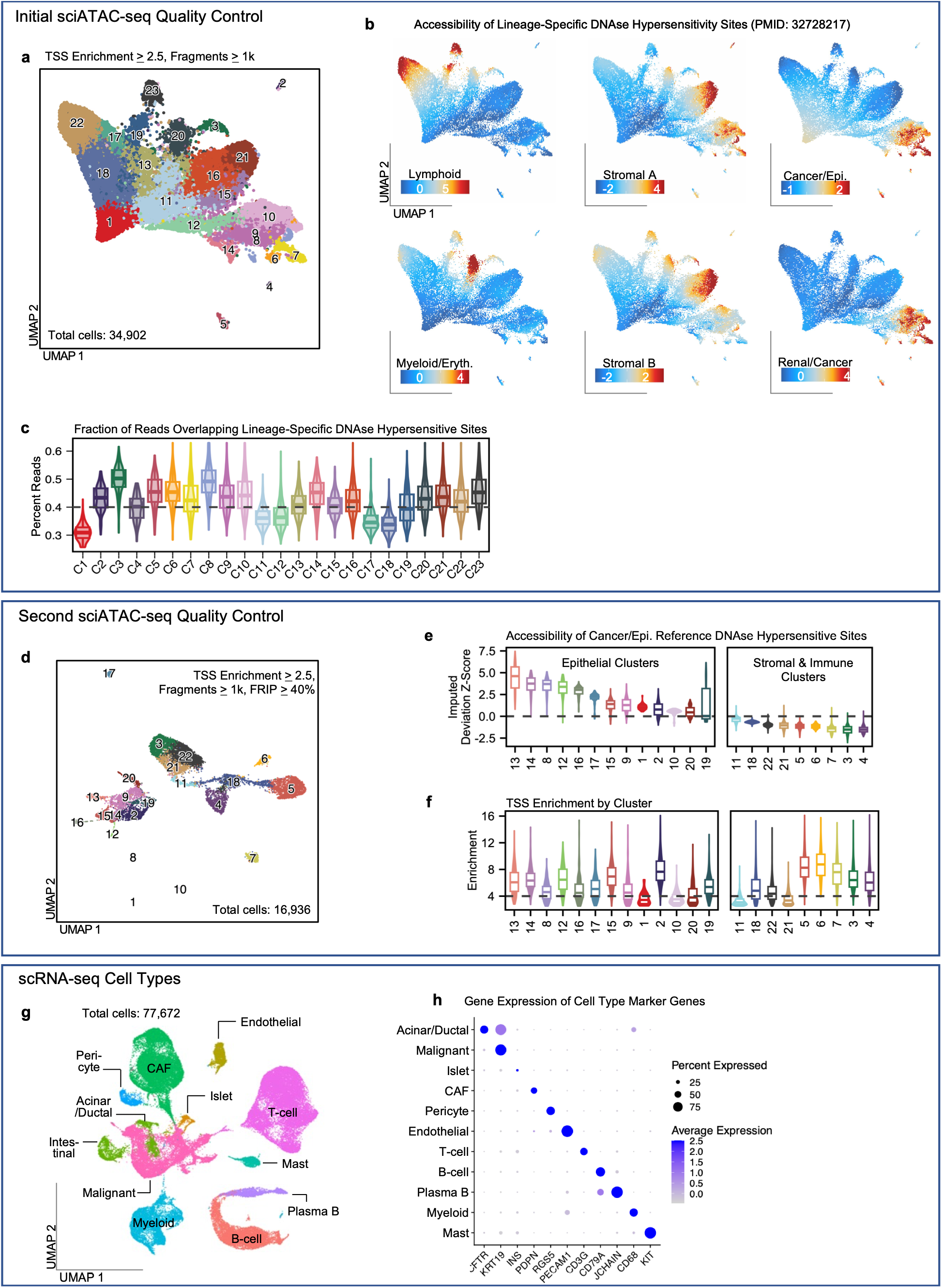
Quality control and cell types in single cell sequencing data. **a)** UMAP embedding and clustering of all single cell epigenomes from all sci-ATAC-seq samples. Cells have at least 2.5 TSS enrichment and 1k fragments. **b)** Per-cell enrichment scores for lineage-associated peak sets by Meuleman W, et al. [72]. Enrichment values are bias-corrected deviation Z-scores calculated by chromVAR. **c)** Per-cell proportion of reads overlapping DNase hypersensitivity sites identified by Meuleman W, et al. [72]. Dotted line indicates the 40% mark. **d)** UMAP embedding and clustering of all single cell epigenomes from all sci-ATAC-seq samples that pass the 40% FRIP cutoff depicted in panel c). **e**) Per-cell enrichment scores for the “Cancer/Epithelial” peak set by Meuleman W, et al. [72]. **f)** Per-cell TSS enrichment score by cluster. Dotted line indicates a 4 TSS enrichment cutoff, which was applied to immune/stromal cells only. **g)** UMAP and clustering results scRNA-seq (n=10 primary tumors). **h)** Expression of cell type marker genes by cell type population shown in panel g).

**Supplementary Figure 2:**
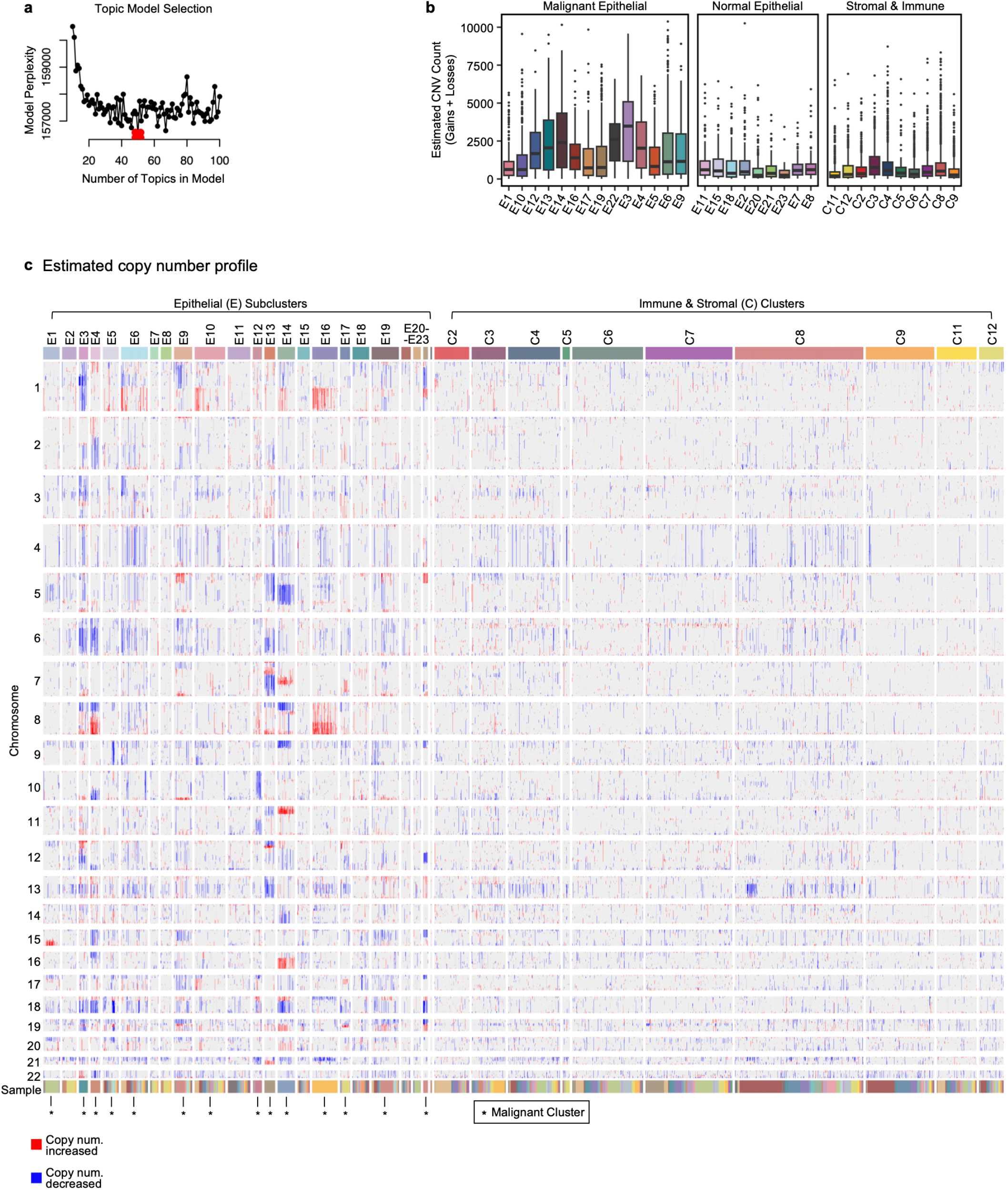
Estimation of copy number in single cell epigenomes. **a)** cisTopic model selection for epithelial/malignant cells from primary tumors and metastases. Red point indicates the model selected in our work. **b)** Distribution of per-cell copy number aberrations (total gains + losses) stratified by cluster. “E” refers to epithelial sub-cluster number (cluster numbers shown in Fig. 1e). “C” cluster numbers refer to the stromal and immune cluster numbers shown in Fig. 1b. **c)** Per-cell copy number changes estimated by epiAneufinder. Rows indicate chromosome number. Columns are divided by single cell cluster; sub-columns within these are individual cells. Asterisks indicate the cluster was deemed malignant.

**Supplementary Figure 3:**
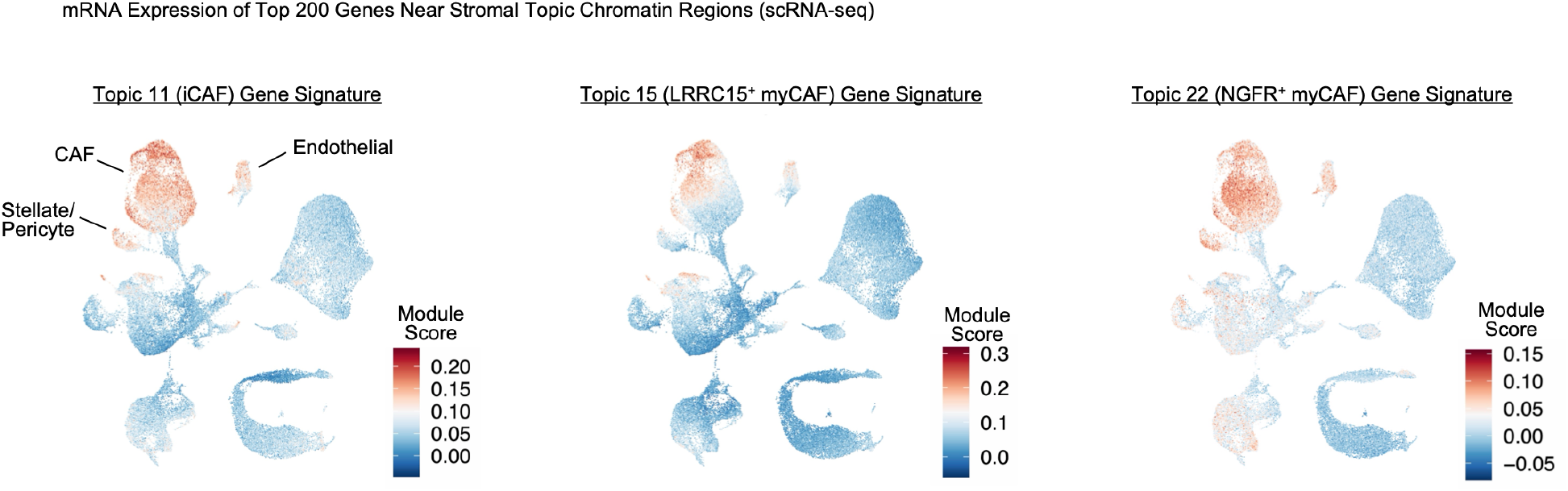
Stromal-specific mRNA expression of gene signatures for epigenetic CAF subtypes. Gene set module scores representing expression of the top 200 genes linked to peaks in Topic 11 (iCAF), Topic 15 (*LRRC15^+^* myCAF), and Topic 22 (*NGFR^+^* myCAF).

**Supplementary Figure 4:**
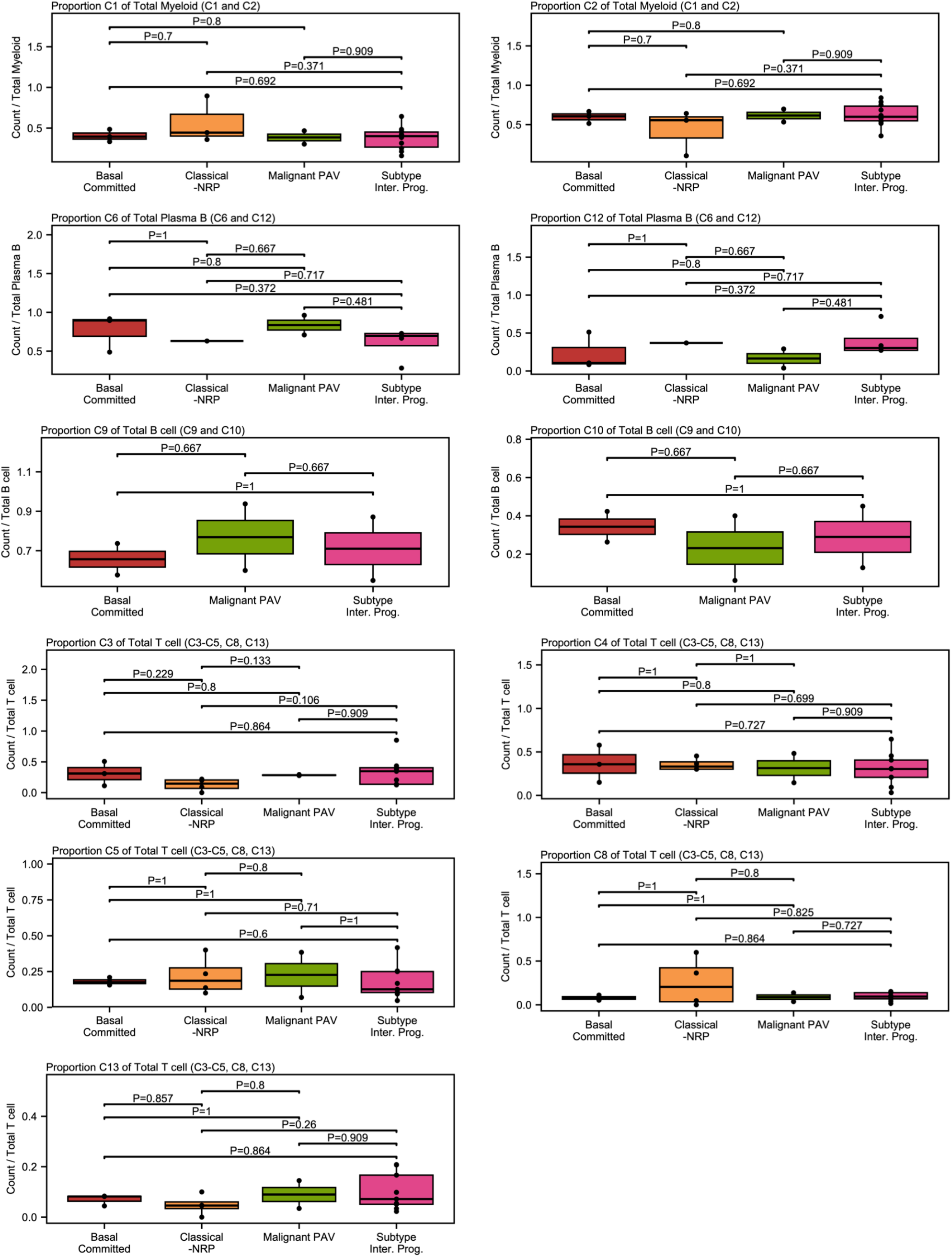
Proportion of immune cell type subpopulations by primary tumor malignant subtype. Proportion of clusters within each immune cell type subpopulation. Proportion is calculated as the total number of cells in the cluster divided by the total number of cells of the relevant cell type. For example, myeloid cluster C1 proportion is calculated as the total C1 cells over the total myeloid cells (C1+C2) for each sample. Samples with fewer than 10 cells of a specific immune cell type are not included in proportion calculations for that cell type. Each point represents a primary tumor; colors indicate the malignant subtype of the primary tumor. Significance was assessed via Wilcox test.

**Supplementary Figure 5:**
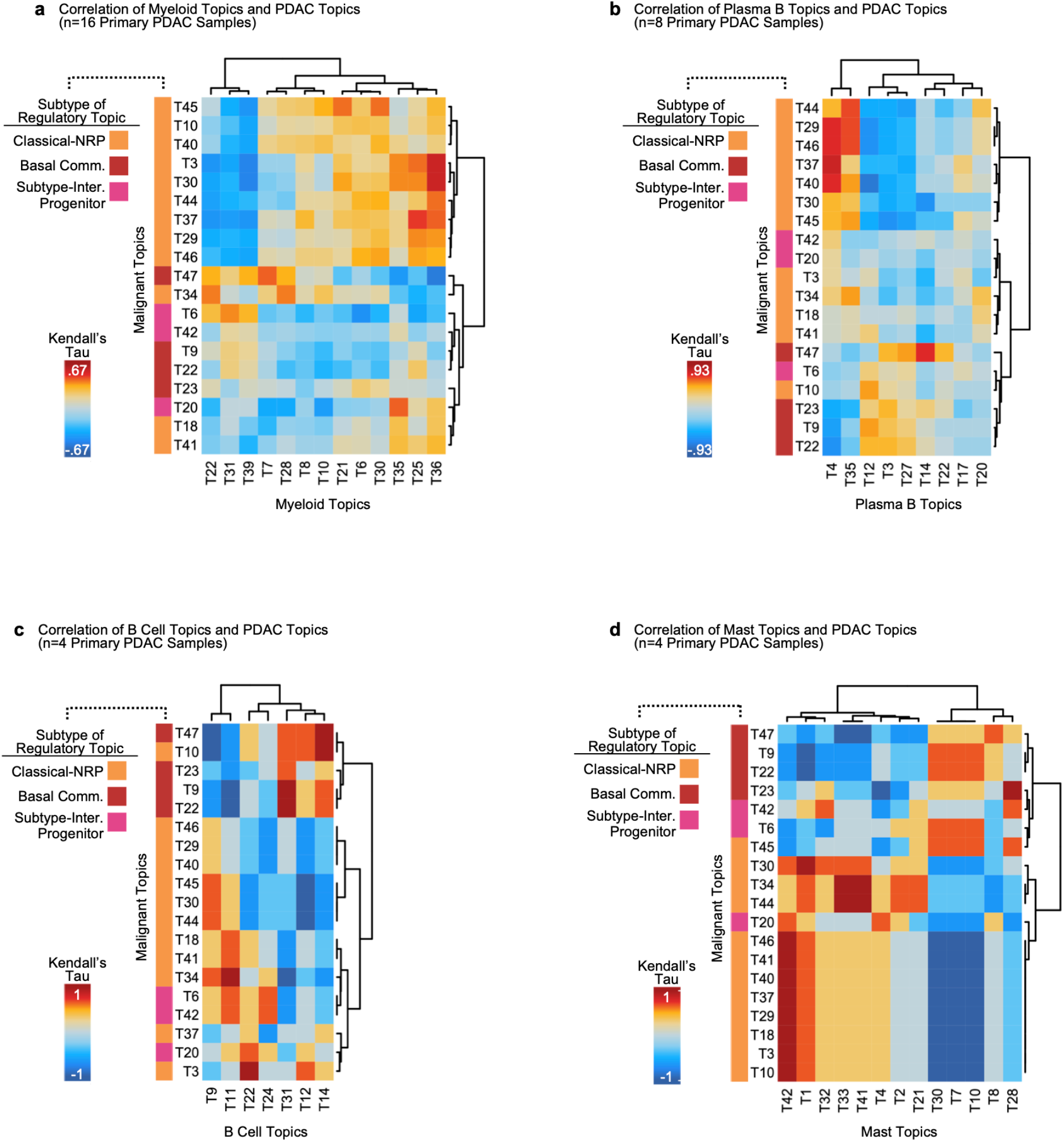
Correlation between immune cell type topics and malignant subtype-associated topics. **a-d)** Sample-level correlation (Kendall’s Tau) between myeloid, plasma B, B cell, and mast-associated immune topics (columns) and malignant subtype-associated topics (rows). Color scale depicts correlation value. Samples with fewer than 10 cells of the relevant immune cell type or fewer than 10 malignant cells are excluded. P-values were FDR-corrected. Asterisks indicate significant results (no significant results were obtained for work presented in this figure).

**Supplementary Figure 6:**
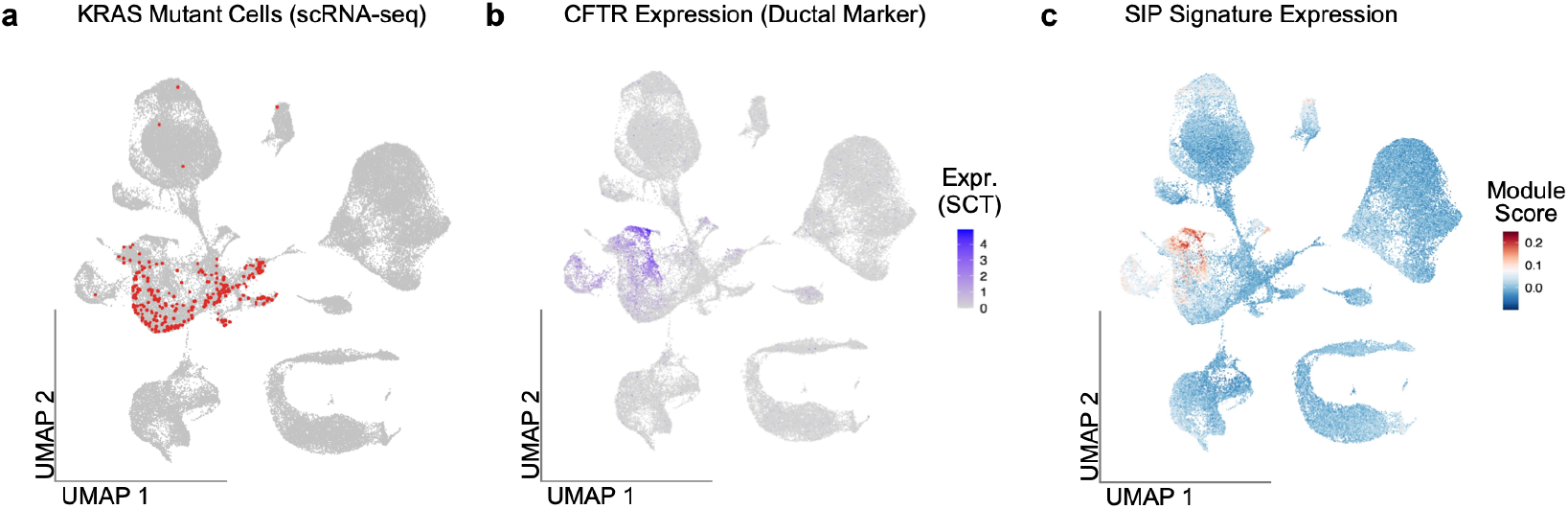
Expression of SIP signature in scRNA-seq data. **a)** scRNA-seq UMAP depicting all cell types from all samples (n=10). Red cells indicate a cell that had one or more *KRAS* mRNA transcripts with a point mutation matching the *KRAS* mutation of the patient the cell originated from. 3 samples were excluded from the survey of *KRAS* mutations surveying due to having no *KRAS* mutation or having unknown *KRAS* status. **b)** Gene expression (SCT values) of the pancreatic ductal marker *CFTR*. **c)** Gene set module scores for the sci-ATAC-seq-derived SIP signature.

